# Targeting Pre-Existing Club-Like Cells in Prostate Cancer Potentiates Androgen Deprivation Therapy

**DOI:** 10.1101/2025.02.13.637653

**Authors:** Manon Baurès, Anne-Sophie Vieira Aleixo, Emeline Pacreau, Aysis Koshy, Vanessa Friedrich, Marc Diedisheim, Martin Raigel, Yichao Hua, Charles Dariane, Florence Boutillon, Lukas Kenner, Jean-Christophe Marine, Gilles Laverny, Daniel Metzger, Florian Rambow, Jacques-Emmanuel Guidotti, Vincent Goffin

**Author notes:** corresponding authors: Vincent Goffin, INEM, 160 rue de vaugirard, 75015, Paris, France and Florian Rambow, Institute for AI in Medicine (IKIM), Department of Applied Computational Cancer Research, Girardetstr. 8, 45131 Essen, Germany. equal contribution (Manon Baurès, Anne-Sophie Vieira Aleixo, Jacques-Emmanuel Guidotti, Vincent Goffin). Current address: ARCHE Core Facility, Biosit UAR 3480 US_S 018, CNRS, Inserm, Université de Rennes, F-35000 Rennes, France.

## Abstract

A critical knowledge gap in prostate cancer research is understanding whether castration-tolerant progenitor-like cells that reside in treatment-naïve tumors play a direct role in therapy resistance and tumor progression. Herein, we reveal that the castration tolerance of LSC^med^ (Lin^−^, Sca-1^+^, CD49f^med^) progenitor cells, the mouse equivalent of human prostatic Club cells, arises not from intrinsic properties, but from significant transcriptional reprogramming. Utilizing single-cell RNA sequencing of LSC^med^ cells isolated from prostate-specific *Pten*-deficient (*Pten^pc-/-^*) mice, we identify the emergence of castration-resistant LSC^med^ cells enriched in stem-like features, driven by the transcription factor FOSL1/AP-1. We demonstrate that cells exhibiting *Pten^pc-/-^* LSC^med^ characteristics are prevalent in aggressive double-negative prostate cancer (DNPC) subtypes recently identified in human castration-resistant prostate cancer (CRPC). Furthermore, our findings show that the dual-targeting agents JQ-1 and CX-6258—focused on FOSL1/AP-1 and PIM kinases, respectively—effectively suppress both the progenitor properties and the growth of mouse and human DNPC surrogates *in vitro* and *in vivo*. Thus, early eradication of castration-tolerant Club-like cells presents a promising therapeutic strategy to mitigate prostate cancer progression toward CRPC.

## Introduction

Prostate Cancer (PCa) is the second most frequent male cancer worldwide and the fifth in terms of death (Bray *et al*, 2018). PCa is androgen-dependent and the gold-standard treatment of advanced PCa is androgen deprivation therapy (ADT), i.e. chemical castration. A clinical response is initially observed in virtually all patients, reflecting the death of androgen-dependent tumoral cells. Within 1 to 3 years, however, castration-resistant PCa (CRPC) develops in most patients, leading to metastatic dissemination to distant sites such as bone and liver (Bubendorf *et al*, 2000). Although CRPC can be further treated with second-generation androgen receptor (AR) pathway inhibitors (ARPi, e.g. abiraterone, enzalutamide, apalutamide, and darolutamide), tumors eventually become resistant within a few months of treatment (Wang *et al*, 2021), which has been linked to the emergence of aggressive forms of CRPC such as mesenchymal stem-like PCa (MSPC) and neuroendocrine PCa (NEPC) (Han *et al*, 2022). JAK/STAT and FGFR signaling (Chan *et al*, 2022), FOSL1/YAP (Tang *et al*, 2022), ONECUT2 (Qian *et al*, 2024), FOXA1 (Eyunni *et al*, 2025) and FOXA2 (Wang *et al*, 2024) were recently identified as drivers of these aggressive forms of CRPC via lineage plasticity, opening new hopes for the treatment of advanced stages of the disease. Therapeutic interventions earlier in the course of the disease to eradicate tumor cells that drive the shift from hormone-sensitive tumors to CRPC may improve the prognosis of PCa patients. Intriguingly, the potential contribution of intrinsic androgen-independent prostatic cells that preexist in naïve prostate tumors is ill-explored.

Club and Hillock cells have been recently identified by single-cell RNA sequencing (scRNAseq) as epithelial cells abundant in the prostatic urethra and collecting ducts, but rare in the healthy human prostate (Henry *et al*, 2018). Club-like cells have also been identified in treatment-naïve primary prostate tumors using scRNAseq and spatial transcriptomics (Chen *et al*, 2021; Kiviaho *et al*, 2024; Song *et al*, 2022). These two cell types exhibit very similar transcriptomic profiles and can be discriminated by the differential expression of lineage markers reflecting a more basal (Hillock) *versus* luminal (Club) phenotype (Henry *et al*., 2018). Club/Hillock cells are enriched in stem/progenitor-like programs and exhibit low androgen signaling, hence they are predicted to be ADT-tolerant (Baures *et al*, 2022a; Henry *et al*., 2018). Accordingly, the transcriptomic signatures of Club/Hillock cells are enriched in the MSPC molecular subtype of PCa (Han *et al*., 2022). MSPC largely overlaps with the stem-cell like (SCL) molecular subtype described by others (Tang *et al*., 2022), both of which are surrogate for double-negative PCa (DNPC) characterized by reduced AR expression signature and no neuroendocrine features (Bluemn *et al*, 2017). Although MSPC is frequently observed at the CRPC stage i.e. post-castration (Han *et al*., 2022), analysis of The Cancer Genome Atlas (TCGA), CPC-GENE (Fraser *et al*, 2017) and DFKZ (Gerhauser *et al*, 2018) datasets reveals that 27-74% of treatment-naïve localized tumors exhibit mixed MSPC and AR-positive PCa (ARPC) features (Han *et al*., 2022). Moreover, up to 11% of the tumors in the early-onset (<55 years of age) DFKZ dataset were identified as MSPC subtype (Han *et al*., 2022). These *de novo* MSPC tumors are associated with enrichment for *PTEN* deletions, more advanced pathology (Gleason score, T/N stages) and premature patient death after treatment compared to ARPC or mixed ARPC/MSPC tumors. The link between biallelic *PTEN* loss and DNPC subtype (57% *versus* 17% frequency in other subtypes, p=0.031) has been recently confirmed (Lundberg *et al*, 2023). Together, these findings converge on the question whether Club cells pre-existing in treatment-naïve tumors may also contribute to therapeutic resistance and tumor progression towards aggressive forms of CRPC. To our knowledge, however, these cells have not been functionally characterized.

The *Pten^pc-/-^* mouse is an appropriate preclinical model to address these key questions as its developing prostatic tumors are highly populated by the mouse equivalent of human prostatic Club/Hillock cells termed LSC^med^ (<u>L</u>in^−^, <u>S</u>ca-1^+^, <u>C</u>D49f<u>^med^</u>) cells (Baures *et al*., 2022a; Baures *et al*, 2022b; Sackmann Sala *et al*, 2017). In this acknowledged mouse model of CRPC, the tumor suppressor gene *Pten* is selectively inactivated in prostatic luminal epithelium, recapitulating one of the most frequent mutations observed in the human disease (Mulholland *et al*, 2011; Wang *et al*, 2003), especially in the DNPC subtype (Han *et al*., 2022; Lundberg *et al*., 2023). According to their low AR-signaling profile, the proliferation rate of *Pten^pc-/-^* LSC^med^ cells is unaffected by castration (Sackmann Sala *et al*., 2017). LSC^med^ cells are enriched in stem/progenitor features (Baures *et al*., 2022a). Accordingly, they exhibit tumorsphere-/organoid-forming capacity *in vitro* (Baures *et al*., 2022b; Sackmann Sala *et al*., 2017) and tumor-initiating capacity both *in situ* (Guo *et al*, 2020) and in graft assays (Sackmann Sala *et al*., 2017). Together, these findings support a key role of these Club-like cells in castration resistance and progression towards CRPC (for a review, Baures *et al*., 2022a).

In this study, we show that *Pten^pc-/-^* LSC^med^ cells are not intrinsically castration-tolerant, but undergo transcriptomic reprogramming after androgen deprivation that enhances their aggressiveness. Using scRNAseq specifically applied to *Pten^pc-/-^*LSC^med^ cells sorted from intact *versus* castrated mice, we show that castration favors the emergence of a LSC^med^ cell subpopulation enriched in stem-like features. We identified the transcription factor FOSL1/AP-1 as the major driver of this transcriptomic phenotype switch. Combined pharmacological inhibition of FOSL1/AP-1 and its transcriptional target PIM1 kinase abrogated the progenitor properties of *Pten^pc-/-^* LSC^med^ cells and markedly reduced the growth of CRPC in castrated *Pten^pc-/-^* mice. In line with the transcriptional resemblance of *Pten^pc-/-^* mouse LSC^med^ cells and the human DNPC subtype, our drug combination also reduced the growth of human MSPC-like PC-3 cells xenografted in castrated host mice. These results suggest that early eradication of castration-tolerant Club-like cells presents a promising therapeutic strategy to mitigate PCa progression toward CRPC.

## Results

### *Pten^pc-/-^* LSC^med^ cells share transcriptomic features with human MSPC/SCL subtypes

LSC^med^ cells isolated from *Pten^pc-/-^* mice exhibit castration tolerance *in vivo* and form organoids in an androgen-independent manner (Baures *et al*., 2022b; Sackmann Sala *et al*., 2017). To investigate whether castration impacts the fate of LSC^med^ cells, we conducted Smart-seq2-based scRNAseq analysis of 384 LSC^med^ sorted cells from dissociated prostates of intact and 2-month-castrated mice (Fig. 1A and Appendix Fig. S1A). After selecting for high-quality single-cell-transcriptomes, we performed dimension reduction using Uniform Manifold Approximation and Projection (UMAP) and annotated the resulting single-cell space according to the corresponding castration profiles (Fig. 1B). The identity of the analyzed cells was assessed by the mRNA expression of the LSC^med^ marker *Krt4* (Sackmann Sala *et al*., 2017) and of the surface markers used in cell sorting (Appendix Fig. S1, B to E). The intrinsically low AR signaling of LSC^med^ cells (Sackmann Sala *et al*., 2017) was further reduced after castration (Fig. 1C). Differential gene expression (DEG) analysis identified 206 genes that exhibited at least a 2-fold (adj. pval<0.001) difference after castration (Table EV1). The 20 most discriminative genes (based on adj. pval and fold-change (FC)) per experimental condition were plotted as heatmap (Fig. 1D). Notably, post-castration, we observed the decreased expression of androgen-regulated genes (*Fxyd3, Mme*) (Mevel *et al*, 2020), and concomitant increased expression of *Krt13,* a Hillock cell marker (Henry *et al*., 2018), and *Krt6b* which is part of the squamous differentiation signature that has been associated with DNPC (Labrecque *et al*, 2019; Lundberg *et al*., 2023). Treatment-induced ARPC to DNPC-squamous conversion has been proposed to be one potential pathway to bypass AR-suppression strategies (Labrecque *et al*., 2019). Finally, various stem-related genes (*Lgr4*, *Klf4*, *Tgfb2, Ly6d*) and inflammatory genes (*Ifi202b, Cxcl5*) were also upregulated. This first experiment shows that *Pten^pc-/-^* LSC^med^ cells are actually sensitive to mouse castration.

**Fig. 1.**
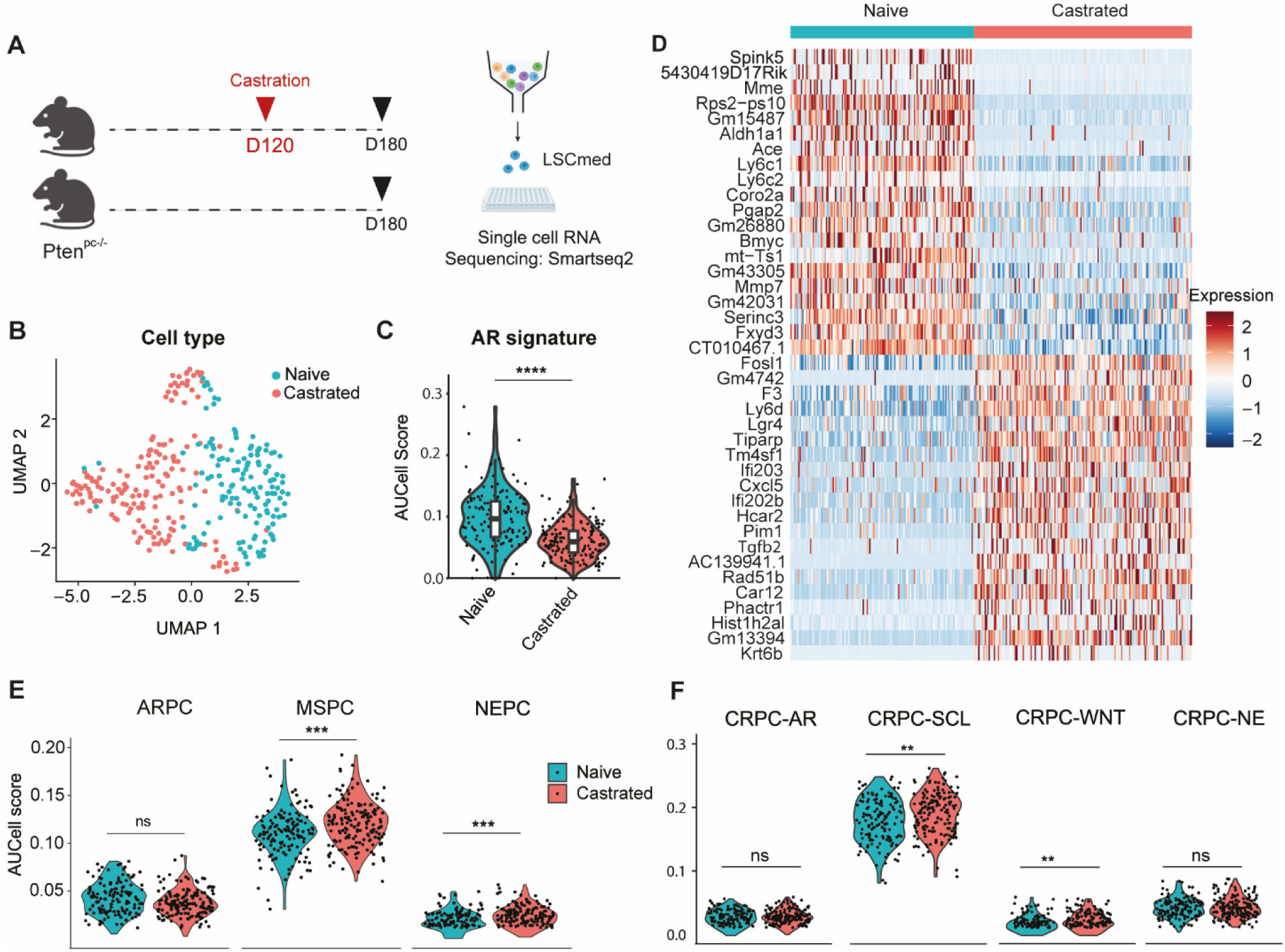
*Pten^pc-/-^*LSC^med^ cells as a robust surrogate of human MSPC subtype. **(A)** Schematic representation of LSC^med^ scRNAseq protocol. Two 6-month-old *Pten^pc-/-^* mice, castrated or not at 4 months of age (Day 120), were used in each group and pooled before scRNAseq analysis. Created in BioRender. BAURES, M. (2024) BioRender.com/f61z820. **(B)** UMAP space of *Pten^pc-/-^* LSC^med^ cells colored according to the castration status. **(C)** AR signature expression intensities (AUCell score) (Sackmann Sala *et al*., 2017) in LSC^med^ cells per *Pten^pc-/-^* mouse, stratified by castration. p=2.18e-15 (Wilcoxon test, Holm-method adjusted). **(D)** The 20 most discriminative (FC and adj. pval) genes of LSC^med^ cells from intact *versus* castrated *Pten^pc-/-^* mice are shown. **(E,F)** The AUCell scores of the MSPC/ARPC/NEPC (Han *et al*., 2022) **(E)** and CRPC-WNT/-SCL/-AR/-NE (Tang *et al*., 2022) **(F)** human CRPC subtype signatures in LSC^med^ cells per *Pten^pc-/-^* mouse, stratified by castration. ns, not significant; **p<0.01; ***p<0.001; ****p<0.0001 (Wilcoxon test, Holm-method adjusted). Exact p-values: p=0.48 (ARPC), p=0.00066 (MSPC), p=0.00017 (NEPC), p=0.57 (CRPC-AR), p=0.0038 (CRPC-SCL), p=0.0058 (CRPC-WNT), p=0.88 (CRPC-NE).

We then compared the signatures of *Pten^pc-/-^* LSC^med^ cells in intact and castrated conditions to recently reported signatures of human PCa subtypes (Table EV2) (Han *et al*., 2022; Tang *et al*., 2022). This analysis revealed that MSPC (Han *et al*., 2022) and its counterpart in the Tang *et al*. study, CRPC-SCL, are markedly enriched in typical LSC^med^ cell genes compared to other PCa subtypes (Fig. 1E and F). The enrichment was further enhanced when using post-castration LSC^med^ cell signatures. This was mainly due to the upregulation of genes such as *Anxa1*, *Tm4sf1*, *Ltbp1*, *Msn*, *Lgals3*, *Arhgdib* (for the MSPC phenotype) and *Plat*, *Sdc4*, *Cstb*, *Cd44*, *Plau*, *Epha2*, *S100a14*, *Pttg1ip* and *Hif1a* (for the SCL phenotype).

Together, these results highlight that *Pten^pc-/-^* LSC^med^ cells are not intrinsically non-responsive to castration and exhibit high molecular similarity with human MSPC/SCL tumors, i.e. DNPC subtypes that are enriched in stemness features and associated with ADT-resistance and metastatic potential.

### Castration strengthens the stem-like characteristics of one *Pten^pc-/-^* LSC^med^ cell subpopulation

To better characterize the impact of castration on *Pten^pc-/-^* LSC^med^ cells, we conducted unsupervised Louvain-clustering of these cells and identified three subpopulations (Fig. 2A and Table EV3). Hereafter these three LSC^med^ clusters are referred to as LSC^med^-0, LSC^med^-1 and LSC^med^-2. The LSC^med^-0 subpopulation, highly predominant before castration, was drastically reduced after castration when LSC^med^-1 subpopulation became predominant (16-fold increased) (Fig. 2B). LSC^med^-2 subpopulation exhibited a more modest increase (3-fold) after castration (Fig. 2B). RT-qPCR analysis of selected subpopulation markers in LSC^med^ cells sorted from intact and 2-month-castrated *Pten^pc-/-^* mice confirmed the opposite regulation of LSC^med^-0 *versus* LSC^med^-1 and LSC^med^-2 genes. Individually, several marker genes of LSC^med^-0 (e.g. *Spink5, Ly6c1*) and LSC^med^- 1 (e.g. *Bcar1, Fosl1, Pim1, F3*) cells exhibited very homogenous response amplitude, which validated the conclusions of scRNAseq experiments (Fig. EV1, A and B). The amplitude of LSC^med^- 2 gene upregulation was more heterogenous (Fig. EV1C). According to the observation reported above for the bulk LSC^med^ cell pool (Fig. 1C), AR signaling was lower in LSC^med^-1 cells, and to a lesser extent, in LSC^med^-2 cells, compared to LSC^med^-0 cells (Fig. 2C).

**Fig. 2.**
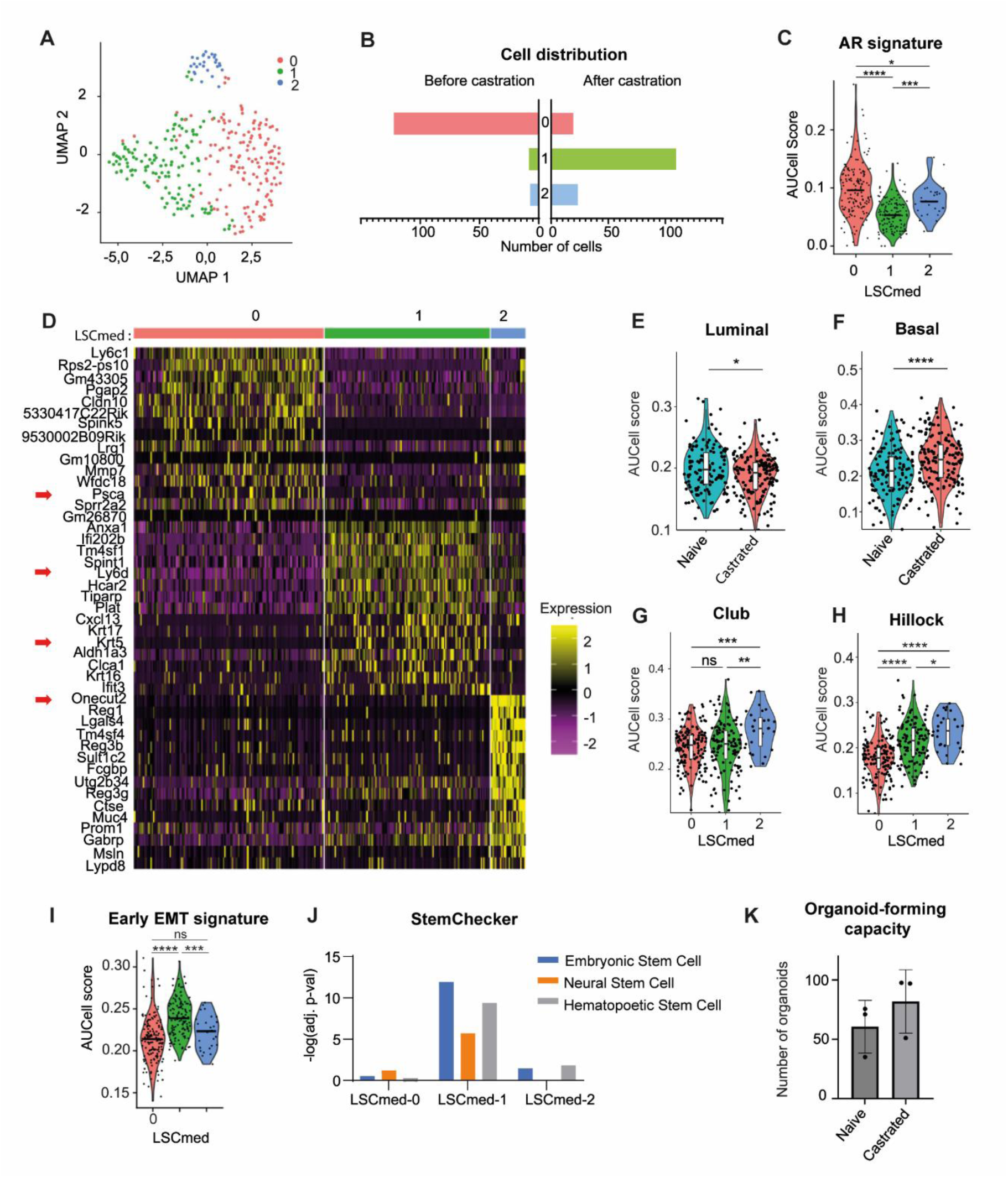
Transcriptomic heterogeneity of *Pten^pc-/-^* LSC^med^ cells. (**A)** The UMAP analysis (unsupervised Louvain-clustering) of *Pten^pc-/-^* LSC^med^ cells analyzed by scRNAseq identified 3 distinct LSC^med^ cell clusters named LSC^med^-0, −1 and −2. (**B)** Distribution of the three LSC^med^ cell subpopulations in prostates from intact *versus* castrated mice (**C)** The AUCell score of the AR signature (Sackmann Sala *et al*., 2017) in the three LSC^med^ cell subpopulations is shown. *p<0.05; ***p<0.001; ****p<0.0001 (Wilcoxon test, Holm-method adjusted). Exact p-values: p=6e-18 (0 *versus* 1), 0.017 (0 *versus* 2), p=0.00017 (1 *versus* 2)**. (D)** Discriminative marker genes (top 15) of the three LSC^med^ cell subpopulations. **(E-I)** Representation of the AUCell scores of the luminal (**E**) and basal (**F**) signatures (Joseph *et al*., 2020b) per castration status, and of the Club (**G**) and Hillock (**H**) signatures (Henry *et al*., 2018) and score of early EMT (**I**) signature (Meyer-Schaller *et al*, 2019) per LSC^med^ cell subpopulations. ns, not significant; *p<0.05; **p<0.01; ***p<0.001; ****p<0.0001 (Wilcoxon test, Holm-method adjusted). Exact p-values: luminal p=0.017; basal p=2.9e-05; Club p=0.49 (0 vs 1), p=0.00085 (0 vs 2), p=0.0072 (1 vs 2); Hillock p=1.3e-10 (0 vs 1), p=7.3e-09 (0 vs 2), p=0.021 (1 vs 2); early EMT p= 5.4 e-14 (0 vs 1), p=0.12 (0 vs 2), p= 0.00076 (1 vs 2). (**J)** StemChecker analysis in each LSC^med^ cell subpopulations (-log (adj. p-values) are represented). (**K)** Organoid-forming capacity of LSC^med^ cells sorted from 10-month-old *Pten^pc-/-^* mice, intact *versus* 2-month-castrated (biological replicates, n = 3 independent experiments). ns, not significant (p= 0,35; unpaired t-test with Welch’s correction).

LSC^med^-0 cells express *Psca*, a well-described androgen-dependent marker in both murine and human luminal progenitors of the prostate (Crowell *et al*, 2019) (red arrow on Fig. 2D and Table EV3). Gene Ontology (GO) enrichment analysis highlighted hallmarks of protein secretion, oxidative phosphorylation, DNA repair and prostate development (Fig. EV1, D and E). With the exception of MYC and E2F targets, most of the pathways in the Hallmark 50 gene dataset were less enriched in LSC^med^-0 cells compared to the two other subpopulations (Fig. EV1D).

LSC^med^-1 cells represent the most amplified subpopulation after castration. It expresses *Ly6d* (red arrow on Fig. 2D and Table EV3), a marker of castration-resistant luminal progenitors that is detected in primary PCa, amplified in mCRPC and associated with biochemical relapse and decreased overall survival (Barros-Silva *et al*, 2018; Steiner *et al*, 2023). Interestingly, we also noted the upregulation of *Krt5* (red arrow on Fig. 2D and Table EV3), associated with an increase in the global basal signature and a slight decrease in the luminal score in response to castration (Fig. 2, E and F), as previously observed in human MSPC (Han *et al*., 2022). This was further confirmed by comparing the transcriptomic signature (top 50 genes) of LSC^med^-1 cells with scRNAseq data of healthy basal and luminal prostatic cells (Data Ref: Crowley *et al*, 2020a; Crowley *et al*, 2020b; Data ref: Joseph *et al*, 2020a; Joseph *et al*, 2020b) (Fig. EV1H). According to the enrichment in basal features, LSC^med^-1 cells showed a stronger Hillock cell profile, while the Club cell profile was unchanged (Fig. 2, G and H). LSC^med^-1 cells were also associated with enrichment in basal breast cancer signature, as well as in other oncogenic pathways (e.g. pancreatic and lung cancer) (Fig. EV1F). Typical features of human MSPC/SCL subtypes (Han *et al*., 2022; Tang *et al*., 2022), including EMT (Fig. 2I), migration and stem signatures (Fig. EV1, D and F), were also enriched in LSC^med^-1 cells. The enrichment of stemness features was further highlighted by StemChecker analysis (Fig. 2J). This analysis allows the association of any input gene list with a curated stemness signature database by calculating enrichment values based on shared genes (hypergeometric test based -log adj. p-value) (Pinto *et al*, 2015). The StemChecker analysis indicated a higher stemness-associated transcriptional identity of LSC^med^-1 cells as suggested by the identification of various stemness-associated genes such as *Klf4* or *Cd44* (Table EV4). Accordingly, GO enrichment analysis highlighted several stemness-associated pathways, e.g. Wnt, Tgfβ, Notch and Hedgehog signaling (green arrows in Fig. EV1D). The trend towards the higher organoid-forming capacity of LSC^med^ cells sorted from the prostates of 2-month-castrated mice (enriched in LSC^med^-1 cells) *versus* intact mice (enriched in LSC^med^-0 cells) agrees with stemness enrichment in LSC^med^ cells post-castration (Fig. 2K and Appendix Fig S2). Together, the molecular features of LSC^med^-1 cells suggest increased aggressiveness of this particular subpopulation amplified following castration.

Finally, the most differentially expressed gene of LSC^med^-2 cells is *Onecut2* (red arrow on Fig. 2D). Onecut2 is known to be a driver of neuroendocrine differentiation in PCa (Chan *et al*., 2022; Guo *et al*, 2019; Qian *et al*., 2024; Rotinen *et al*, 2018). In LSC^med^-2 cells, *Onecut2* expression was associated with low enrichment (AUCell score values <0.05) of a neuroendocrine signature (Han *et al*., 2022) (Fig. EV1I), but typical neuroendocrine markers such as Chromogranin A and Synaptophysin were not detected in LSC^med^-2 cell transcriptome. In contrast to LSC^med^-1 cells, LSC^med^-2 cells were not enriched in stemness-associated programs (Fig. 2J) and exhibited less basal features (Fig. EV1H). GO enrichment analysis highlighted the increased oxidative stress and hypoxia in LSC^med^-2 cells. Of note, the hypoxia-associated enzyme transglutaminase 2 (*Tgm2*), which is a marker of LSC^med^-2 cells (Table EV3), was recently identified as a marker of malignant progression (Abu El Maaty *et al*, 2022). This suggests that the LSC^med^-2 subpopulation could have the potential to promote cancer progression, which could be facilitated by various oncogenic pathways associated with tumor aggressiveness (Fig. EV1G).

Together, these results demonstrate that castration of *Pten^pc-/-^*mice promotes the emergence of LSC^med^ cells presenting with a transcriptomic profile enriched in basal, EMT and stemness features and lineage plasticity drivers. All these features are known to correlate with cancer cell aggressiveness.

### Transcriptomic plasticity of *Pten^pc-/-^* LSC^med^ cells as an adaptive mechanism to castration

The transcriptomic analysis described above identified three LSC^med^ cell subpopulations whose ratios considerably evolve in response to castration (Fig. 2B). We hypothesized that this could result from a phenomenon of i) positive selection, i.e. robust proliferation of LSC^med^-1 cells possibly associated to death of LSC^med^-0 cells, and/or ii) cell plasticity, i.e. a transcriptional switch from LSC^med^-0 to LSC^med^-1 and LSC^med^-2 profiles (Fig. 3A). To address these hypotheses, we first quantified Ki-67 positive epithelial cells in various fields of prostates harvested from *Pten^pc-/-^* mice sacrificed prior to castration or at 5, 21 or 60 days after castration (Appendix Fig. S3A). Confirming our previous findings (Sackmann Sala *et al*., 2017), cell proliferation was not altered following castration (Fig. 3, B and C). The pattern of dead cells (TUNEL assay) was focal and the global level remained particularly low (∼2.5%), with no significant variation after castration (Fig. 3D and Appendix Fig. S3, B and C). In contrast, we showed by RT-qPCR that the expression of DEGs associated with each LSC^med^ cell subpopulation quickly evolved after castration. Downregulation of LSC^med^-0 markers (e.g. *Hoxb13, Spink5*) and concomitant upregulation of LSC^med^-1 markers (e.g. *Ifi202b, Tm4sf1*) were detected as early as 5 days after castration (Fig. 3, E and F). Although the pattern of LSC^med^-0 (down) *versus* LSC^med^-1 (up) markers displayed a clear time-dependent response, the variability observed between the 3 biological replicates was consistent with a dynamic process. For LSC^med^-2 cell markers, fold-changes were overall more modest (with the exception of *Reg1*), suggesting that a very low number of cells exhibited these features within 21 days post-castration. The rapid emergence of LSC^med^-1 cells after castration was confirmed by immunohistochemistry showing the upregulation of CD44, identified as a marker of this subpopulation (Fig. EV2, A and B, and Table EV3).

**Fig. 3.**
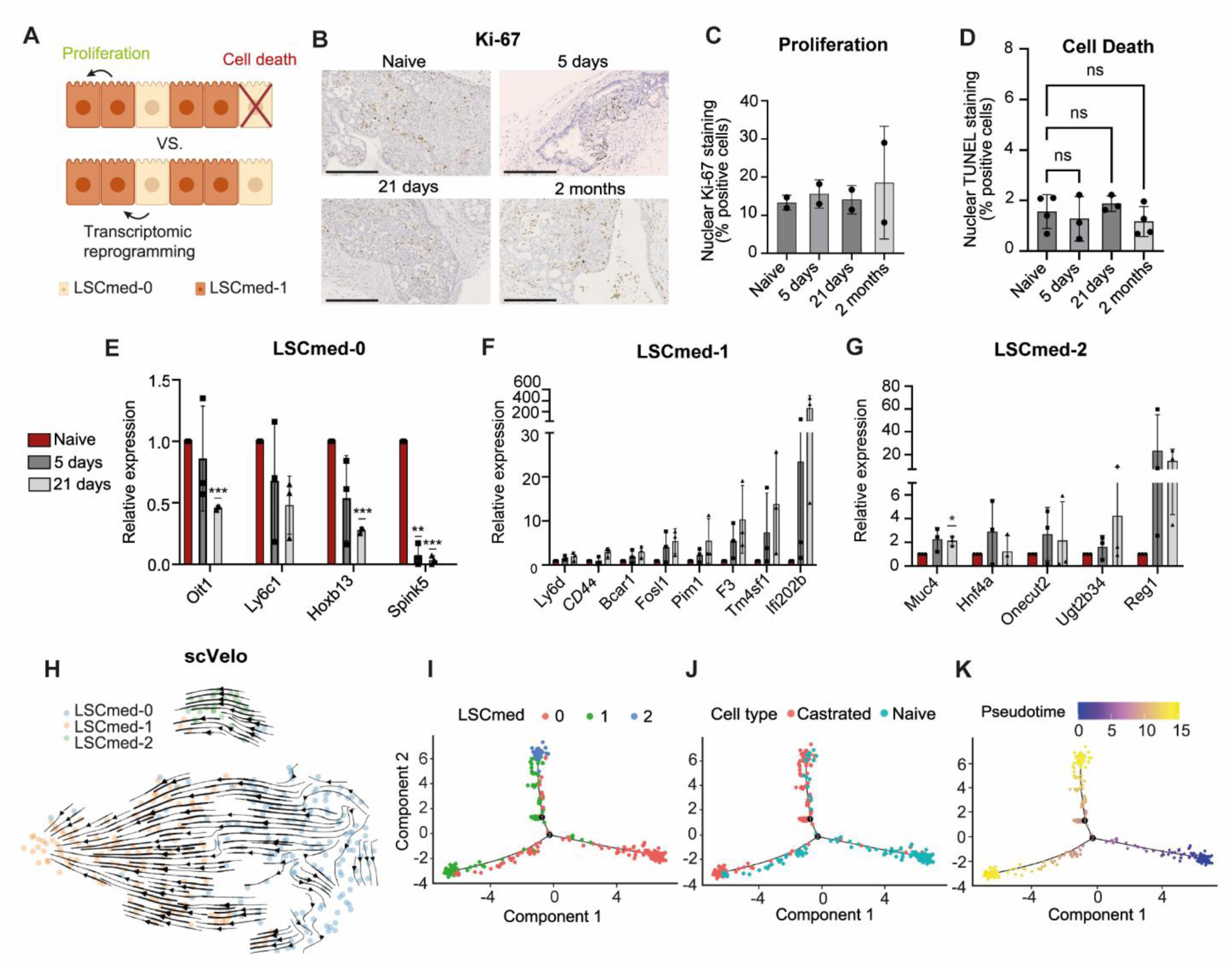
Transcriptional plasticity dynamics of *Pten^pc-/-^* LSC^med^ cells after castration. **(A)** Schematic representation of positive selection (upper panel) *versus* transcriptional reprogramming (lower panel) as two non-mutually exclusive mechanisms that could promote amplification of LSC^med^-1 and LSC^med^-2 at the expense of LSC^med^-0 cell subpopulations after castration (see text for explanations). Created BioRender. BAURES, M. (2024) BioRender.com/k26b175. **(B,C)** Images **(B)** and proportion of Ki67-positive cells **(C)** quantified from immunohistochemistry (scale bar: 250 μm) in prostates from intact *Pten^pc-/-^*mice *versus* 5 days, 21 days and 2 months after castration, 2 mice per condition, 5 to 10 fields counted per animal (See Appendix Table S3A for total cell counts per animal). **(D)** TUNEL-positive cells in prostates from intact *Pten^pc-/-^* mice *versus* 5 days, 21 days and 2 months after castration, 3 to 4 mice per condition, 9 to 23 fields counted per animal (See Appendix Table S3B for total cell counts per animal). ns, not significant (Welch’s ANOVA with Brown-Forsythe correction, Dunnett’s T3 post hoc test). Exact p-value: p= 0, 0,9469 (naive vs 5 days), p= 0,8017 (naive vs 21 days), p= 0,7613 (naive vs 2 months). **(E-G)** The expression of selected marker genes of LSC^med^-0 **(E)**, LSC^med^-1 **(F)** and LSC^med^-2 **(G)** cells was measured by RT-qPCR in bulk LSC^med^ cells sorted from intact *Pten^pc-/-^* mice and 5 and 21 days after castration. N=3 independent experiments, each biological replicate corresponds to one to three pooled animals for a total of n=5 (naive) and n=7 (5 and 21 days). Data are normalized to the expression in intact mice. *p<0.05; **p<0.01; ***p<0.001 (unpaired t-test with Welch’s correction). Exact p-values are reported in Appendix Table S1. **(H)**. RNA velocity analysis (scVelo) in *Pten^pc-/-^* LSC^med^ cells analyzed by scRNAseq, based on the inference of directed dynamic information by leveraging splicing kinetics. **(I-K)** Pseudotime-ordering analysis (Monocle): projection of cells per cluster **(I),** castration status **(J)** and pseudotime **(K)**.

The concomitance of early transcriptional changes with the unaltered rates of cell proliferation and cell death favors a transcriptional reprogramming trajectory from LSC^med^-0 towards LSC^med^-1 cells shortly after castration, while emergence of LSC^med^-2 cells may occur at a later stage. RNA velocity analysis (scVelo), based on the inference of directed dynamic information by leveraging splicing kinetics, predicted LSC^med^-0 cells as a likely root cell population (Fig. 3H). This prediction was further supported by pseudotime-ordering analysis (Monocle) (Fig. 3, I to K). The transcriptional heterogeneity of LSC^med^-0 and LSC^med^-1 cells was evident from the presence of “intermediate” cells distributed along the lower trajectory before castration (Fig. 3, I to K). In contrast, the grouping of all LSC^med^-2 cells at the upper branch extremity suggests higher transcriptional homogeneity. Trajectory analyses predicted that post-castration LSC^med^-1 cells evolve from pre-castration LSC^med^-0 cells, while post-castration LSC^med^-2 cells may originate from LSC^med^-0 and/or LSC^med^- 1 cells.

These results collectively support the hypothesis that cellular plasticity underlies an adaptive resistance mechanism of LSC^med^ cells, which is initiated shortly after castration.

### AP-1 complex is a major regulator of *Pten^pc-/-^* LSC^med^ cells emerging post-castration

We next performed gene regulatory inference using SCENIC (Aibar *et al*, 2017) to identify transcriptional upstream regulators of the three LSC^med^ cell subpopulations. This analysis predicted *Hoxb13* as a major regulator of LSC^med^-0 cells (Fig. 4A and Fig. EV3A). *Hoxb13* was described as key driver of the prostatic luminal cell differentiation (Huang *et al*, 2007). Other highlighted transcription factors regulating LSC^med^-0 cells included *Ddit3*, *Bmyc*, *Nr2f2*, *Atf3*, *Grhl2*, and *Ehf* (Fig. 4A and Fig. EV3A). *DDIT3* was recently shown to be upregulated in CRPC patients and proposed to be associated with progression to CRPC (Jung *et al*, 2024). EHF is a transcription factor of the ETS family that controls epithelial cell differentiation and its loss promotes prostate tumors enriched in EMT signature and mesenchymal/stem-like features (Albino *et al*, 2012). Finally, while *Bmyc* may be correlated to the upregulation of MYC targets (Fig. EV1D), *Nr2f2* has been shown to cooperate with *Pten* deletion to promote malignant progression (Qin *et al*, 2013).

**Fig. 4.**
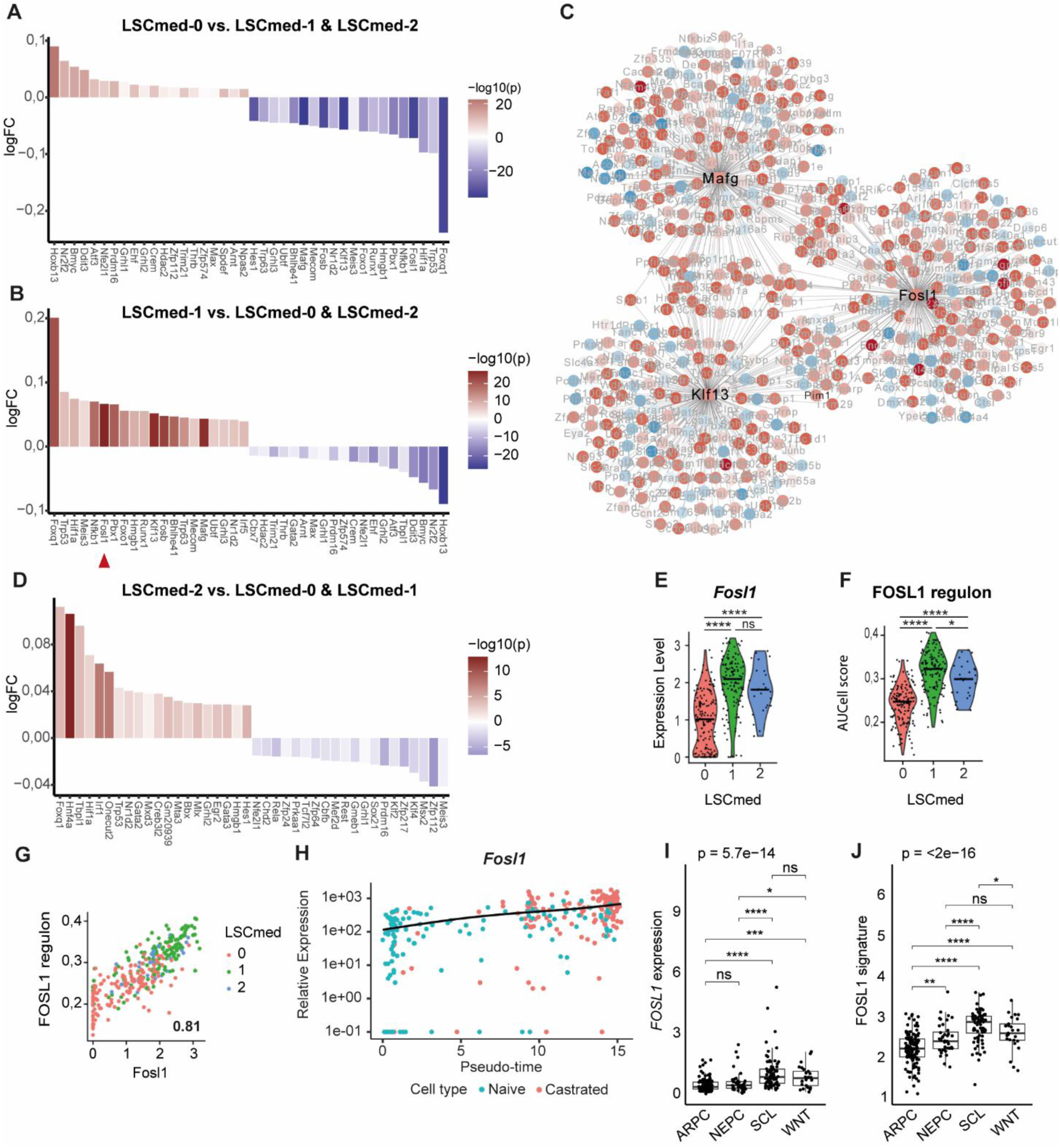
FOSL1/AP-1 complex is a major transcriptional hub for *Pten^pc-/-^* LSC^med^ cells emerging post-castration. **(A to D)** Identification of the main upstream regulators of the three LSC^med^ cell subpopulations by SCENIC analysis. The top-20 (fold change) activated (red) and inactivated (blue) transcription factors in LSC^med^-0 **(A)**, LSC^med^-1 **(B)** et LSC^med^-2 **(D)** cells are represented as waterfall plots. The red arrow in B identifies *Fosl1*. The transcription factor network of the 3 most pertinent regulons of LSC^med^-1 cells is represented in **C** (see Fig. EV3 for LSC^med^-0 and LSC^med^-2 cell transcription factor networks). **(E,F)** *Fosl1* expression (**E**) and AUCell Score of FOSL1 regulon (**F**) in the three LSC^med^ cell subpopulations. ns, not significant; *p<0.05; ****p<0.0001 (Wilcoxon test, Holm-method adjusted). Exact p-values: *Fosl1*: p=3.7e-21 (0 vs 1), 1.3e-06 (0 vs 2), p=0.11 (1 vs 2); Fosl1 regulon: p=4.8e-26 (0 vs 1), 1e-07 (0 vs 2) and p=0.031 (1 vs 2). **(G)** Correlation of *Fosl1* expression and FOSL1 regulon in the three LSC^med^ cell subpopulations. **(H)** Projection of *Fosl1* expression during pseudotime (pval = 5.01e-21, qval = 6.37e-19). **(I,J)** Expression of *FOSL1* **(I)** and FOSL1 regulon **(J)** in SU2C patient cohort (Data Ref: Abida *et al*., 2019a), according to the tumor molecular subtype (n=125 ARPC, n=40 NEPC, n=76 SCL, n=25 WNT). ns, not significant; **p<0.01; ***p<0.001; ****p<0.0001 (Kruskal-Wallis with Holm-corrected Wilcoxon post-hoc tests). See Appendix Table S1 for exact p-values.

Remarkably, the transcription factor profile of LSC^med^-1 cells (Fig. 4B) was the near-perfect inverse mirror image of that of LSC^med^-0 cells (Fig. 4A). This observation further supports that castration triggers cell plasticity of LSC^med^-0 towards LSC^med^-1 cells by switching on/off selected transcription factors. We identified *Fosl1* as the most pertinent (based on -log10(pval)) regulator of the LSC^med^-1 cell signature (Fig. 4B). FOSL1 is a transcription factor that acts in the AP-1 complex mainly as a heterodimer involving Jun proteins (c-Jun, JunB, JunD) (Eferl & Wagner, 2003). Other typical transcription factors of LSC^med^-1 cells included *Mafg, Klf13*, *Foxq1*, *Nfκb1*, *Pbx1*, *Ets1* and *Fosb* (Fig. 4B). Of note, both *Mafg* and *Klf13* share several common target genes with FOSL1 (Fig. 4C).

Finally, we identified *Hnf4a* as a master regulator of LSC^med^-2 cell signature (Fig. 4D and Fig. S3B). HNF4a is a nuclear receptor known for its role as a tumor suppressor in the prostate by its ability to promote senescence in prostatic cells (Wang *et al*, 2020). HNF4a shares several target genes with Oncecut2 (Fig. EV3B), suggesting a potential role of HNF4a in the cellular plasticity leading to the LSC^med^-2 signature. *Nr1d2*, another transcriptional regulator promoting lineage plasticity towards NEPC (He *et al*, 2021), was also identified as a top gene of LSC^med^-2 cells (Fig. 4D). The expression of *Gata2* and *Gata3* (Fig. 4D) suggests that LSC^med^-2 cells present a certain degree of luminal differentiation (Xiao *et al*, 2016).

Although *Fosl1* was identified as the top gene of LSC^med^-1 cells (Fig. 4B), it was expressed at similar level in LSC^med^-2 cells, and at consistent, though lower, level in LSC^med^-0 cells (Fig. 4E). The FOSL1-regulated network expectedly followed the same pattern (Fig. 4, F and G). FOSL1/AP-1 is known to be involved in various processes including stemness, EMT and cell plasticity (Eferl & Wagner, 2003; Feldker *et al*, 2020; Marques *et al*, 2021). In the mouse prostate, FOSL1/AP-1 was recently identified as one of the actors of differentiated (androgen-dependent) luminal cell plasticity in both healthy (Kirk *et al*, 2024) and cancer (Tang *et al*., 2022) contexts. To investigate further the potential role of FOSL1/AP-1 complex in Club-like cell plasticity upon castration, we plotted the *Fosl1* expression in function of pseudotime. As illustrated in Fig. 4H, we observed that *Fosl1* expression progressively increased post-castration. The upregulation of FOSL1 protein after castration was confirmed by immunohistochemistry (Fig. EV2, C and D). As frequently observed in cancer cells (Sobolev *et al*, 2022; Song *et al*, 2006; Taha *et al*, 2023), both cytoplasmic and nuclear FOSL1 staining were observed, the latter being mainly detected post-castration. According to previous reports (Casalino *et al*, 2022), castration-triggered FOSL1 activation may contribute to the upregulation of its own gene expression. The analysis of sorted LSC^med^ cells by immunoblotting further confirmed that castration upregulates FOSL1 expression in this particular cell population (Fig. EV2E). Together, these data suggest that the FOSL1/AP-1 regulon is a main driver of the post-castration transcriptional switch of LSC^med^ cells. Other AP-1 factors could contribute as their transcriptional networks were also highlighted by SCENIC analysis, albeit at lower significance (*Atf3* in LSC^med^-0 cells, *Fosl2* and *Fosb* in LSC^med^-1 cells, *Fos* in LSC^med^-2 cells) (Fig. 4, A, B and D, and Table EV3). Strikingly, in human PCa, FOSL1 and its regulon are particularly associated with the MSPC/SCL subtype (Fig. 4, I and J) (Data Ref: Abida *et al*, 2019a; Abida *et al*, 2019b). In fact, we noticed that the majority of AP-1 complex members (*FOSL1*, *FOSL2*, *FOS*, *JUN*, *JUNB*, *ATF3*, *BATF*) are expressed at higher levels in MSPC/SCL than in the ARPC tumors (Fig. 4, I and J, and Fig. EV4, A to F). This data further highlights the relevance of *Pten^pc-/-^* LSC^med^ cells as a surrogate of the aggressive DNPC subtype.

Together, these analyses show that members of the AP-1 complex, and in particular FOSL1, are predicted to be major drivers of the transcriptional switch occurring post-castration in LSC^med^/Club-like cells.

### Pharmacological targeting of LSC^med^ and Club/Hillock-like cancer cells *in vitro*

We next used *Pten^pc-/-^* LSC^med^ cells as a DNPC surrogate to identify relevant therapeutic targets for this incurable cancer subtype. Considering that LSC^med^-1 cells are the most amplified after castration (Fig. 2B), the least androgen-sensitive (Fig. 2C) and the most enriched in stemness features (Fig. 2J), we reasoned that targeting this particular cell subpopulation was the most relevant strategy to deviate LSC^med^ cells from progression towards CRPC in our mouse model. This hypothesis was strengthened by the observation that, in the recent DARANA study (Linder *et al*, 2022a; Data ref: Linder *et al*, 2022b), 3-month enzalutamide treatment of naïve high risk PCa patients led to a marked enrichment of prostate tumors in LSC^med^-1 signature at the expense of LSC^med^-0 signature (Fig. EV5A), mimicking what we observed in *Pten^pc-/-^* mice after castration (Fig. 2B). Based on the results shown in the previous section, FOSL1/AP-1 was identified as the most obvious therapeutic target to interfere with LSC^med^-1 cell emergence. The high expression of various members of the AP-1 family, including *Fosl1*, in the two other LSC^med^ subpopulations (Fig. EV4, G to L) suggested a wide spectrum of targeted LSC^med^ cells. The markedly increased expression of *FOSL1* and FOSL1 target genes after enzalutamide treatment of PCa patients in the DARANA study (Linder *et al*., 2022a; Data ref: Linder *et al*., 2022b) further supports the relevance of FOSL1 as a therapeutic target in the human disease (Fig EV5, B and C).

We noticed that *Pim1* is part of the *Fosl1* gene network in the mouse (Fig. 4C). PIM1 is a member of the PIM serine/threonine kinase family that phosphorylates substrates controlling various tumorigenic phenotypes, including cell survival and proliferation (Rout *et al*, 2024; Wang *et al*, 2001). In PCa, it was shown to promote cell invasion and migration (Rebello *et al*, 2016; Santio *et al*, 2015) and ligand-independent androgen receptor phosphorylation associated with hormone-refractory PCa (Ha *et al*, 2013). PIM1 pharmacological targeting was shown to reduce prostate tumor growth in mouse and human preclinical settings (Hu *et al*, 2009; Rebello *et al*., 2016). We found that *Pim1* is drastically enriched in *Pten^pc-/-^*LSC^med^-1 and LSC^med^-2 cell subpopulations which are predominant post-castration (Fig. 5, A and B).

**Fig. 5.**
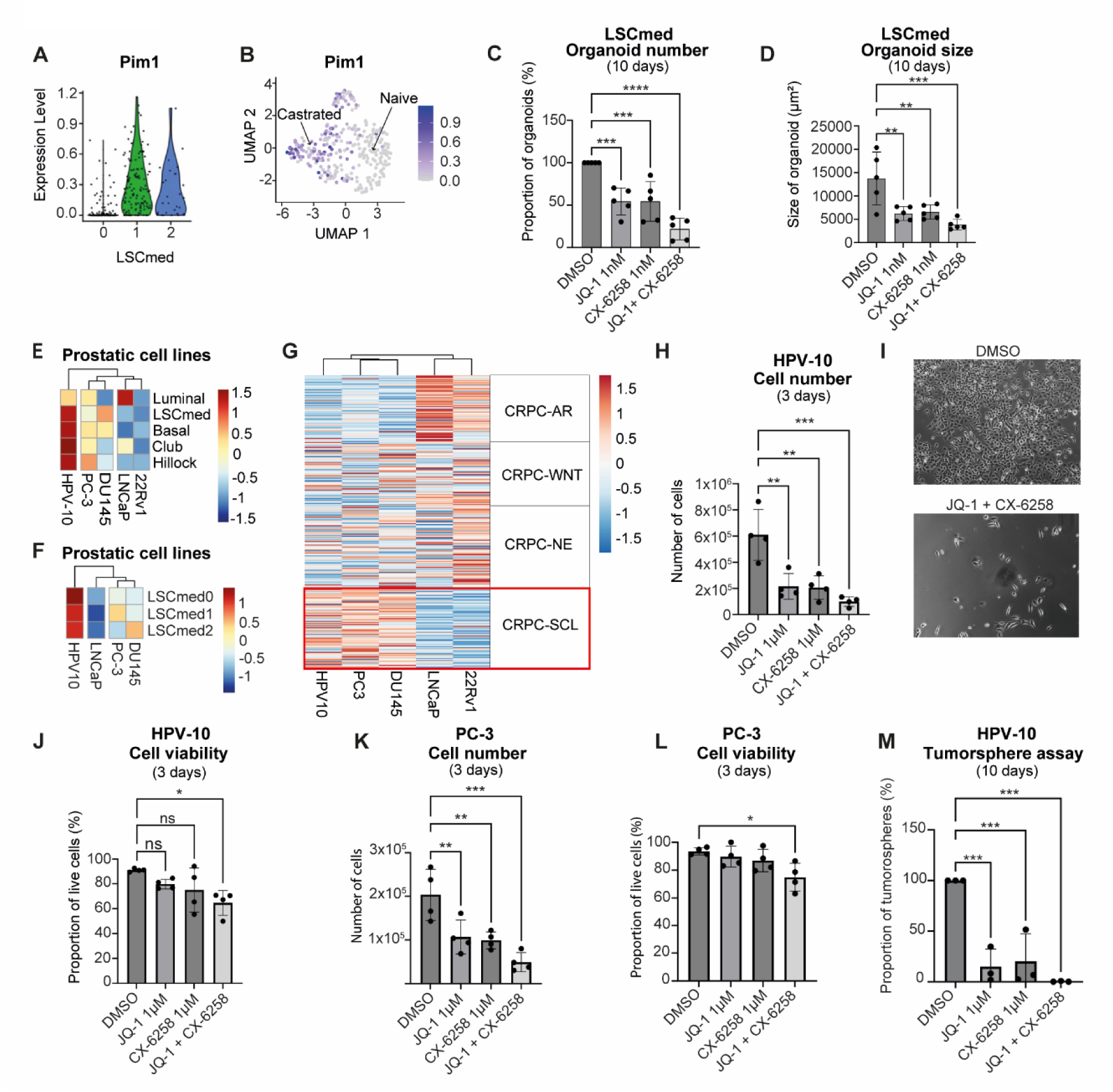
Targeting FOSL1/AP-1 and Pim family prevents the growth of LSC^med^ and human Club-like cells *in vitro*. **(A,B)** Expression per *Pten^pc-/-^* LSC^med^ cell subpopulation **(A)** and UMAP projection **(B)** of *Pim1*. **(C,D)** Number **(C)** and size **(D)** of organoids generated from sorted *Pten^pc-/-^* LSC^med^ cells after 10 days of culture in medium containing DMSO or 1 nM JQ-1 and 1 nM CX-6258, alone or combined, as indicated. Data are normalized to the control (DMSO) condition (biological replicates, n=5 independent experiments). **p<0.01; ***p<0.001; ****p<0.0001 (One-way ANOVA and Dunnett *post hoc* test). Exact p-values are reported in Appendix Table S1. **(E,F)** Enrichment of various epithelial cell signatures **(E)** and LSC^med^ subpopulation signatures **(F)** in HPV-10, PC-3, DU145, LNCaP and 22Rv1 cell lines (Data Ref: Wang *et al*., 2007a). **(G)** Enrichment of CRPC-AR, CRPC -WNT, CRPC -NE and CRPC -SCL human tumoral subtypes signatures (Table EV2) in HPV-10, PC-3, DU145, LNCaP and 22Rv1 cell lines. **(H)** Human HPV-10 cells were treated for 72 hours with 1 µM JQ-1 and 1 µM CX-6258, alone or combined (as indicated), then the number of adherent cells was counted (biological replicates, n = 4 independent experiments). The data are normalized to the control condition (DMSO). **p<0.01; ***p<0.001 (One-way ANOVA and Dunnett *post hoc* test). Exact p-values are reported in Appendix Table S1. **(I)** Images of HPV-10 cells after 72h of treatment. **(J)** The viability of HPV-10 cells (adherent + in suspension) was determined by trypan blue staining (biological replicates, n=4 independent experiments). ns, not significant; *p<0.05 (One-way ANOVA and Dunnett *post hoc* test). Exact p-values are reported in Appendix Table S1. **(K,L)** Same as H and J with PC-3 cells (biological replicates, n=4 independent experiments). ns, not significant; *p<0.05 **p<0.01; ***p<0.001 (One-way ANOVA and Dunnett *post hoc* test). Exact p-values are reported in Appendix Table S1. **(M)** Tumorsphere-forming capacity of HPV-10 cells in medium containing DMSO or 1µM JQ-1 and 1 µM CX-6258, alone or combined as indicated. Data are normalized to the DMSO condition (biological replicates, n=3 independent experiments). ***p<0.001 (One-way ANOVA and Dunnett *post hoc* test). Exact p-values are reported in Appendix Table S1.

Of the two other paralogs of the Pim kinase family, *Pim3*, but not *Pim2*, is highly expressed in all *Pten^pc-/-^* LSC^med^ cell subpopulations (Fig. EV4, M and N). Of note, *PIM1* expression was also increased by enzalutamide treatment in naïve PCa patients albeit less markedly than *FOSL1* (Fig. EV5D).

We tested JQ-1 and CX-6258 as pharmacological inhibitors of these two selected targets in functional assays. JQ-1 is a global BET/AP-1 inhibitor that has been shown to mainly target FOSL1 in lung adenocarcinomas (Casalino *et al*., 2022; Lockwood *et al*, 2012) and to phenocopy *FOSL1* knockdown in other types of tumors (Baker *et al*, 2015; Bid *et al*, 2016). Importantly, JQ-1 had been validated in preclinical *in vivo* settings prior to this study (Shimamura *et al*, 2013; Shu *et al*, 2016). CX-6258 is a pan-PIM kinase inhibitor (Haddach *et al*, 2012), which prevents compensation by any other expressed Pim paralog (e.g. Pim3 in our case) as reported in other studies (Mikkers *et al*, 2002; van der Lugt *et al*, 1995). CX-6258 was recently shown to interfere with PCa progression in preclinical models (Rebello *et al*., 2016).

We used the organoid assay to determine the functional impact of these pharmacological inhibitors on *Pten^pc-/-^* LSC^med^ cell properties. We first determined the dose-response of each drug on the organoid-forming capacity of these cells. Both JQ-1 and CX-6258 were very active in the nanomolar range, indicating high sensitivity of LSC^med^ cells to these drugs (Fig. EV6, A and B). At 1 nM concentration, each drug decreased by half both the number and the size of the organoids generated from sorted *Pten^pc-/-^* LSC^med^ cells, indicating inhibition of their progenitor and proliferation capacities, respectively. These effects were further potentiated when both drugs were combined (Fig. 5, C and D). These data indicate that targeting FOSL1/AP-1 and/or PIM kinases markedly alters the growth and progenitor properties of *Pten^pc-/-^*LSC^med^ cells.

We then aimed to validate this therapeutic approach using human Club/Hillock cancer cells. In contrast to *Pten^pc-/-^* mouse LSC^med^ cells, there are currently no established procedures to sort, culture and expand Club/Hillock cells from fresh specimens of human prostate cancer. Therefore, we sought a relevant cell line matching the Club/Hillock phenotype. By comparing human PCa cell line signatures (Data Ref: Wang *et al*, 2007a; Wang *et al*, 2007b) to mouse LSC^med^ (Sackmann Sala *et al*., 2017) and human Club, Hillock (Henry *et al*., 2018), basal and luminal (Pitzen *et al*, 2025) cell signatures (Table EV5), we found that HPV-10 cells are highly enriched in LSC^med^/Club/Hillock features (Fig. 5E). In comparison, androgen-independent PC-3 and DU145 cell lines exhibited an intermediate enrichment, while androgen-dependent LNCaP and 22Rv1, classically described as luminal prostate cancer cells, were the least enriched in LSC^med^/Club/Hillock features among the cell panel analyzed (Fig. 5E).

The HPV-10 cell line is not commonly used in the field. In contrast to PC-3 and DU145 cells, which were derived from PCa metastatic sites, HPV-10 cells were isolated from a high-grade primary prostate adenocarcinoma. They grow in culture in the absence of androgens, which indicates their intrinsic castration tolerance. Accordingly, no *PSCA* expression was detected by RT-qPCR (Appendix Fig. S4A). Otherwise, we confirmed the expression of various LSC^med^ markers (e.g. *KRT4*, *KRT7*) in HPV-10 cells compared to the luminal LNCaP cell line (Appendix Fig. S4A). Immunofluorescence analysis showed the co-expression of KRT4 (Club cell marker) and KRT13 (Hillock cell marker) (Appendix Fig. S4B), identifying HPV-10 as a mixed Club/Hillock cell line. This is reminiscent of murine LSC^med^ cells that also match both Club and Hillock signatures (Baures *et al*., 2022a; Baures *et al*., 2022b). HPV-10 cells are particularly enriched in the LSC^med^-0 signature (Fig. 5F), and they express *FOSL1* and *PIM1* at a much higher level than LNCaP cells, further supporting their Club-like cell profile (Fig. EV6C and Appendix Fig. S4C). While HPV-10 cells are not tumorigenic, the accumulation of chromosomal alterations, including amplification of c-MYC, characteristic of tumorigenic PC-3 cells, makes them tumorigenic (Hukku *et al*, 2000). Altogether these data identify the HPV-10 cell line as a relevant model of Club/Hillock PCa cells preexisting in prostate tumors, i.e. at a less aggressive stage than metastatic PC-3 cells that are exclusively enriched in LSC^med^-1 signature (Fig. 5F). Finally, HPV-10 cells, but not LNCaP and 22Rv cells, expressed the genes of the CRPC-SCL signature to a similar degree as PC-3 and DU145 cells, previously identified as MSPC/SCL models (Han *et al*., 2022; Tang *et al*., 2022) (Fig. 5G and Table EV2). This identifies HPV-10 cells as another surrogate of DNPC subtype.

To determine the effects of JQ-1 and CX-6258 on human Club/Hillock-like cells, we first performed dose-response assays using the HPV-10 cell line (Fig. EV6D). For both drugs, we observed a significant inhibition of the number of adherent cells in the micromolar range, indicating lower drug sensitivity compared to *Pten*^pc-/-^ LSC^med^ cells (Fig. EV6, A and B). Based on that data, we used 1 µM of each drug for subsequent experiments. As observed above with *Pten*^pc-/-^ LSC^med^ cells, the combination of JQ-1 and CX-6258 potentiated the inhibition of HPV-10 cell growth in two-dimension cultures (Fig. 5, H and I). The moderate (15-25%) ratio of dead cells at 72h of treatment (Fig. 5J) suggested that the drugs mainly prevented cell proliferation, as reported in former studies using JQ-1 (Li *et al*, 2019; Zhang *et al*, 2020; Zhang *et al*, 2021). A similar pattern of drug response was observed with PC-3 cells (Fig. 5, K and L). In addition, we found that the tumorsphere formation by HPV-10 cells was significantly altered by the drugs, again with almost a total inhibition when using the drug combination (Fig. 5M). This assay could not be performed with PC-3 cells as these cells failed to generate tumorspheres in our hands.

To confirm the FOSL1-dependence of the effects observed with JQ-1, we looked for an alternative FOSL1 inhibitor. T5224 is an AP-1 inhibitor that was recently shown to directly bind to FOSL1 (Zaman *et al*, 2024) and to inhibit tumorigenesis with low/moderate potency in various cancer models (Kamide *et al*, 2016; Tang *et al*., 2022). A PROTAC version of T5224 was reported to selectively degrade FOSL1 in head and neck squamous cell carcinoma, leading to increased potency compared to the parental compound (Zaman *et al*., 2024). Treatment of PC-3 cells with T5224-PROTAC led to similar effects compared to JQ-1 treatment, alone or combined to CX-6258 (Fig. EV6, E and F). HPV-10 cells exhibited even higher drug sensitivity compared to PC-3 cells (Fig. EV6, G and H). These data strengthen the role of FOSL1 in Club-like cell growth and survival.

To definitely establish this conclusion, we aimed to silence *FOSL1* expression in PC-3 and HPV-10 cells. HPV-10 cells proved extremely sensitive to lipofectamine, which *per se* was detrimental to cell survival at lipofectamine concentrations required for siRNA entry into cells. In contrast, efficient *FOSL1* knock-down in PC-3 cells was obtained with three different siRNAs (Fig. EV6I). Using the most potent of them (siFOSL1(1)), we observed that *FOSL1* silencing significantly altered the growth and viability of PC-3 cells (Fig. EV6, I to L). These effects were quantitatively similar to those obtained with 15 µM T5224-PROTAC (Fig. EV6, E and F).

Together, these results indicate that FOSL1 is a relevant target, and JQ-1/CX-6258 combination an effective therapy, to abolish the progenitor and growth properties of mouse and human Club-like prostate cancer cells *in vitro*.

### Targeting the FOSL1/AP-1 and Pim family markedly reduces CRPC growth *in vivo*

We then aimed to validate this drug combination strategy in a CRPC context *in vivo*. Based on the progressive upregulation of *Fosl1* and *Pim1* expression in LSC^med^ cells soon after *Pten^pc-/-^* mouse castration (Fig. 3F, Fig. EV1B and Fig. EV2, C to E), we reasoned that castration-induced reprogramming of LSC^med^-0 towards LSC^med^-1 transcriptional profiles should provide a window of opportunity for combination therapy involving JQ-1 and CX-6258.

Two groups of experimental mice (n=8 each) were castrated and after 4-day recovery, we started treatment with JQ-1 (daily) and CX-6258 (bi-weekly) *versus* vehicle (Fig. 6A). After 28 days, mice were euthanized, then prostates were harvested, weighed, and processed for histopathological analyses or functional assays. The kidneys, lungs, and livers of five animals from both the control and treated groups were also harvested and examined by two pathologists blinded to group assignments to assess drug toxicity. Only minor pathological lesions were observed (Appendix Table S2), with no significant differences between the control and treated groups (Appendix Fig. S5, A and B). We concluded that these are incidental lesions, and the administered drug combination had no toxic effect at that dosage. The absence of treatment-induced cell death as determined by the TUNEL assay fully supported this conclusion (Appendix Fig. S5C).

**Fig. 6.**
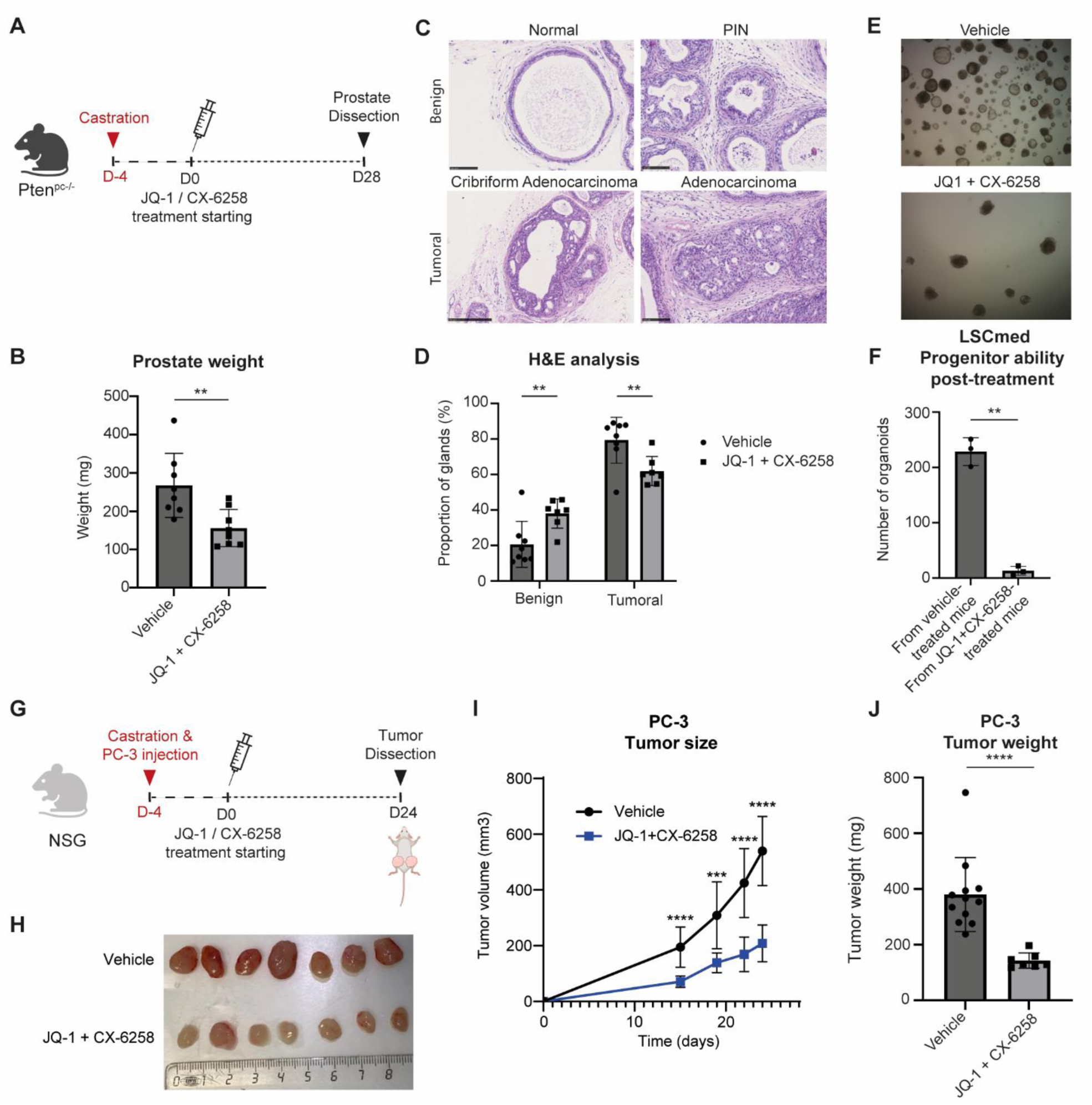
Combined JQ-1 and CX-6258 treatment reduce CRPC growth and aggressiveness *in vivo*. **(A)**. Timeline representation of the experimental protocol used for *Pten^pc-/-^*mouse castration and pharmacological treatments. Castrated mice received vehicle (n=8) or daily JQ-1 intraperitoneal injection and bi-weekly CX-6258 oral administration (n=8) during 28 days. (**B**) After 28 days of treatment, prostates were micro-dissected and weighed. **p<0.01 (unpaired two-tailed *t* test with Welch’s correction). Exact p-value: p=0.0075. **(C)** For all mice, prostate histological structures were classified as benign (normal glands, PINs) or tumoral (cribriform, adenocarcinoma) as described in the Materials and Methods (scale bar = 100 μm). **(D)** Quantification of the effects of pharmacological treatments on prostate histopathology illustrated in panel C. **p<0.01 (unpaired two-tailed *t* test with Welch’s correction). Exact p-values: p=0.0087 (benign), p=0.0087 (tumoral). **(E,F)** After 28 days of treatment, LSC^med^ cells were sorted from the prostate of vehicle- or drug-treated animals and cultured in organoid medium in the absence of pharmacological inhibitors. Image were taken (**E**) and the number of organoids was measured (**F**) after 10 days of culture (biological replicates, n=3 experiments). ***p<0.001 (unpaired two-tailed *t* test with Welch’s correction). Exact p-value: p=0.0023. (**G)**. Timeline representation of the experimental protocol used for PC-3 cell xenograft assay and pharmacological treatments. 1.10^6^ PC-3 cells were injected into each flank of castrated immunodeficient NSG mice. Mice were randomized to receive vehicle (n=12) or daily JQ-1 intraperitoneal injection and bi-weekly CX-6258 oral administration (n=7) during 24 days. **(H-J)** During the protocol, tumor size was measured manually at days 15, 19, 22 and 24 of treatment (**I**). **p<0.01; ***p<0.001; ****p<0.0001 (unpaired two-tailed *t* test with Welch’s correction comparing vehicle and JQ-1 + CX-6258 at each time). Exact p-values: p<0.0001 (D15), p=0.0004 (D19, p<0.0001 (D22), p<0.0001 (D24). At sacrifice (D24), tumors were micro-dissected (**H**) and weighed (**J**). ***p<0.001 (unpaired two-tailed *t* test with Welch’s correction). P-value: p<0.0001. Panels A and G were created in BioRender. BAURES, M. (2024) BioRender.com/p76n322

We observed that the prostate weight in the drug-treated group was reduced by 41% compared to the vehicle-treated group (Fig. 6B). This was associated with a decreased proportion of advanced histologically-classified structures at the benefit of benign phenotypes (i.e. normal glands and prostate intraepithelial neoplasia [PINs]) (Fig. 6, C and D). In parallel to these studies, LSC^med^ cells sorted from a set of fresh prostates of vehicle- or drug-treated mice (each n=4) were cultured in organoid medium for 10 days in the absence of JQ-1 and CX-6258. Remarkably, we observed that LSC^med^ cells sorted from drug-treated *Pten^pc-/-^* mice had virtually lost any capacity to generate organoids compared to the vehicle-treated group (Fig. 6, E and F).

To determine whether JQ-1/CX-6258 treatment interfered with the emergence of the LSC^med^-1 cell subpopulation, as expected, the whole transcriptome of EpCAM+ cells in prostate sections from vehicle-*versus* drug-treated castrated *Pten^pc-/-^* mice was analyzed by Digital Spatial Profiling. Prostate sections from naïve (non-castrated) *Pten^pc-/-^* mice were used as a control in this experiment. This analysis confirmed the marked enrichment of the LSC^med^-1 signature score in castrated *Pten^pc-/-^* mice (Fig. EV7A). No significant variation in the LSC^med^-0 and LSC^med^-2 signature scores could be identified in the glands of the 3 cohorts analyzed (Appendix Fig. S7), possibly reflecting the lower sensitivity of this bulk transcriptomic profiling technology compared to scRNAseq (Fig. 2B). Importantly, the LSC^med^-1 signature score was reduced by JQ-1/CX-6258 treatment, although it did not reach the score of naïve mice (Fig. EV7A). In total, 21 genes were significantly deregulated (p-value <0.05 and a log2 fold-change >0.5) in castrated mice after the treatment (*C3, Lgals3bp, Srrm2, Srgn, Ifi203, Ifi27l2a, Ifitm3, Ctsl, Oas2, Apobr, Pglyrp1, Hsd11b2, Atp2a3, H2-D1, H2-Q6, Basp1, S100a6, Cd74, H2-K1, Psmb8, Hspa1a*).

All top genes affected by the treatment in both LSC^med^-1 (*C3, Cd74, Hspa1a, Ifi203, Ifi27l2a, Ifitm3*) and LSC^med^-2 (*S100a6, Srgn*) signatures were downregulated (Fig. EV7B). These genes are mainly involved in inflammatory pathways, such as genes involved in the complement system (i.e. C3), antigen presentation (e.g. Cd74), and IFN-α and -γ pathways (e.g. Ifitm3). Note that for some transcripts, the treatment normalized their levels (*Hspa1a*, *S100a6* and *Srgn*). Taken together, these results confirm that the LSC^med^-1 signature is amplified in *Pten^pc-/-^* mice after castration and that a FOSL1/PIM kinase inhibitor targets this cell subpopulation.

Next, we set up a xenograft assay to assess the *in vivo* efficacy of our therapeutic strategy on human DNPC cells. We used the PC-3 cell line as HPV-10 cells have been characterized as non-tumorigenic *in vivo* (Hukku *et al*., 2000; Weijerman *et al*, 1994). Immunodeficient NSG mice were castrated and PC-3 cells were then injected subcutaneously into both flanks of the animals before the start of treatment four days later (Fig. 6G). Follow-up of tumor volume showed that tumor growth was markedly delayed in the drug-treated group compared to the vehicle-treated group (Fig. 6, H and I, and Appendix Fig. S6). On day 24, tumors were excised and weighed (Fig. 6J). Both the volume (- 62%) and the weight (- 59%) of tumors were drastically reduced in the drug-treated group.

Together, these data demonstrate that pharmacological targeting of FOSL1/AP-1 and PIM kinases using the combination of JQ-1 and CX-6258 inhibitors markedly reduces the growth *in vivo* of androgen-independent MSPC/SCL-like cells in a castration context.

## Discussion

A critical knowledge gap in prostate cancer research is to understand whether castration-tolerant progenitor-like cells that reside in treatment-naive tumors play a direct role in castration resistance, making them relevant therapeutic targets. This study uncovers the unexpected finding that pre-existing Club-like cells are not intrinsically non-responsive to androgen deprivation but actually respond to castration by a transcriptional switch that increases their aggressiveness. This transcriptional plasticity is mainly orchestrated by FOSL1/AP-1 complex. This finding opens new therapeutic avenues to eradicate treatment-resistant PCa cells early in the course of the disease for suppressing progression towards CRPC. Supporting this promising therapeutic approach, combined pharmacological targeting of FOSL1/AP-1 and PIM kinases markedly reduced the *in vivo* tumor growth capacities of preclinical DNPC models, which was correlated to the suppression of their progenitor properties *in vitro*.

The concept of pre-existing CRPC cells in treatment-naïve tumors was pioneered by the group of Risbridger using patient-derived xenografts of treatment-naïve early-stage tumors (Toivanen *et al*, 2013). These authors showed that, four weeks after mouse castration, residual tumor foci contained stem-like tumor cells able to regenerate proliferating tumors upon androgen replenishment. However, the co-expression of stem-like (NANOG, ALDH1, CD44) and differentiated (NKX3.1) cell markers cannot rule out that these CRPC cells actually arose from castration-induced luminal cell plasticity, a mechanism of castration resistance recently documented in various mouse models (Chan *et al*., 2022; Karthaus *et al*, 2020; Kirk *et al*., 2024). More recently, the existence of stem-like/EMT-enriched CRPC cells in treatment-naïve prostate tumors has been revealed by scRNAseq approaches. These cells have been correlated with biochemical recurrence (rising PSA levels) and distant metastasis (Cheng *et al*, 2022). Other authors (Chen *et al*., 2021) identified one basal/intermediate epithelial cell population (referred to as cluster 10) that we subsequently showed to be enriched in typical Club/Hillock cell markers (Baures *et al*., 2022b). This cluster was identified in each individual patient, albeit at highly variable ratio (1% to 25% of the epithelial cell pool). Song and colleagues classified PCa-associated Club-like cells into six transcriptomic states, the largest of which exhibited the highest luminal and AR signaling characteristics (Song *et al*., 2022). This is also what we observed for the *Pten*^pc-/-^ mouse LSC^med^-0 subpopulation predominant before castration. All these scRNAseq studies corroborate the report by Han and colleagues who showed on large cohorts that the MSPC subtype, enriched in Club cell features, is present *de novo* in a significant proportion of treatment-naïve PCa (Han *et al*., 2022).

According to the molecular similarity of mouse LSC^med^ and human Club/Hillock cells, we established that *Pten*^pc-/-^ mouse LSC^med^ cells are a robust surrogate of human DNPC/MSPC/SCL subtype. Based on this finding, the *Pten*^pc-/-^ mouse model was attractive as it shows a highly consistent enrichment of LSC^med^ cells in tumors prior to castration (Sackmann Sala *et al*., 2017), mimicking in an amplified way human *de novo* MSPCs. LSC^med^-like cells have been classically considered as an end-stage of androgen-independent cells. Our study shows that this is not the case as naïve *Pten*^pc-/-^ LSC^med^ cells, despite of their intrinsically low androgen-signaling, are sensitive to castration and respond to androgen deprivation by a transcriptional switch. Before castration, *Pten*^pc-/-^ LSC^med^ cells mainly exhibit the LSC^med^-0 profile enriched in developmental signaling pathways, as previously observed for wild-type LSC^med^-like luminal progenitor cells that participate in the early steps of prostate morphogenesis (Baures *et al*., 2022a; Mevel *et al*., 2020). In contrast, post-castration *Pten*^pc-/-^ LSC^med^ cells mainly exhibited the LSC^med^-1 profile, which is comparatively enriched in stemness and basal/Hillock features. This finding echoes a recent report showing that DNPC is enriched in cells exhibiting a mixed basal, Club, and Hillock identity (Pitzen *et al*., 2025). This profile, which is typical of the human MSPC subtype resistant to ARPIs (Han *et al*., 2022), further increases the transcriptomic similarity between post-castration LSC^med^ cells and the MSPC/SCL subtypes. The acquisition of an intermediate basal/luminal phenotype by cell plasticity has also been reported in triple-negative breast cancer and it was shown to promote resistance to chemotherapy (Marsolier *et al*, 2022). In the prostate context, the acquisition of a basal/stem cell signature identifies aggressive PCa phenotypes (Smith *et al*, 2015). This signature was also enriched in hormone-sensitive metastases compared to organ-confined prostate adenocarcinomas, suggesting that the enrichment of basal features parallels PCa progression. Therefore, the acquisition of basal features by LSC^med^-like cells post-castration could be pivotal for their aggressiveness and the emergence of MSPC-CRPC tumors in response to treatment.

We identified FOSL1/AP-1 as a major driver of LSC^med^ cell plasticity in response to castration. While the AP-1 complex is known to mediate cellular stress, the detection of FOSL1 upregulation by various technological approaches using tissue samples immediately processed after animal sacrifice (e.g. IHC, RT-qPCR) eliminates any technical bias due to cellular stress potentially induced by methodologies involving cellular dissociation of prostatic tissue (e.g. cell sorting, scRNAseq). The AP-1 complex, and in particular FOSL1, is involved in the development of many adenocarcinomas (Casalino *et al*., 2022). Acting as a transcription factor, facilitator of chromatin accessibility/opening and/or super enhancer (Bi *et al*, 2020; Dong *et al*, 2021; Kadur Lakshminarasimha Murthy *et al*, 2022), FOSL1 regulates various tumor-associated processes e.g. cell proliferation (Zanconato *et al*, 2015), EMT (Feldker *et al*., 2020; Marques *et al*., 2021), invasion and metastases (Iskit *et al*, 2015), and stemness (Marques *et al*., 2021). Via its association with partners such as YAP and TAZ (Feldker *et al*., 2020; Zanconato *et al*., 2015), FOSL1 also promotes cell plasticity towards increased stemness states (Marques *et al*., 2021) and resistance to treatment (Bi *et al*., 2020). FOSL1 has been recently shown to be involved in the reprogramming of mouse prostatic luminal cells towards a castration-tolerant progenitor-like state following androgen depletion (Tang *et al*., 2022). We here identify FOSL1 as a master regulator of LSC^med^-1 cells, i.e. the predominant LSC^med^ population after castration. The consistent expression levels of *Fosl1* and of various members of the AP-1 complex in both LSC^med^-1 and LSC^med^-2 cells strengthens the key role of this transcriptional complex in the molecular switch of pre-existing LSC^med^ cells in response to castration. In humans, *FOSL1* has been identified as the top key transcription factor in CRPC-SCL (Tang *et al*., 2022). It is positively associated to the chromatin remodeling that promotes CRPC-SCL subtype in castrated context by enhancing CRPC-SCL-associated enhancer accessibility, in cooperation with the YAP-TAZ complex (Tang *et al*., 2022). As observed for post-castration LSC^med^ cells, human CRPC-SCL is enriched in various AP-1 family members. Furthermore, we found that both *FOSL1* and FOSL1-target genes are upregulated by enzalutamide treatment of naïve PCa patients (Fig. EV5 and Linder *et al*., 2022a). Together, these data support the key role of this transcriptional complex in the adaptation of treatment-naïve PCa luminal cells – both differentiated (Tang *et al*., 2022) and progenitor-like (this study) – to androgen deprivation.

Based on our findings, we reasoned that inhibition FOSL1/AP-1 pathway should suppress the castration-driven mechanisms of resistance of LSC^med^ progenitor cells promoting CRPC. This therapeutic strategy was strengthened by coupling pharmacological inhibition of PIM kinase, previously shown to promote cell stemness (Gao *et al*, 2019; Jimenez-Garcia *et al*, 2016) and to affect PCa growth (Hu *et al*., 2009; Rebello *et al*., 2016). The combination of castration and JQ- 1/CX-6258 treatment of *Pten*^pc-/-^ mice markedly reduced *in situ* CRPC growth and histopathology after only 28 days compared to castration alone. Similar tumor growth reduction was observed with xenografted PC-3 cells. One remarkable effect of this pharmacological treatment was to abrogate the progenitor properties of *Pten*^pc-/-^ LSC^med^ cells as determined from their organoid/tumorsphere-forming capacity. Remarkably, this effect was also observed with LSC^med^ cells harvested from JQ- 1/CX-6258-treated *Pten*^pc-/-^ mice, without adding drugs in the organoid assay (Fig. 6J). Such a radical inhibition of their ability to form organoids cannot be explained solely by the reduction in the LSC^med^-1 signature revealed by spatial transcriptomic analysis. Although it is at present unknown whether this treatment suppress the progenitor properties of some LSC^med^ cells, or eradicate the progenitor cells *per se,* our data establish a correlation between *in situ* tumor growth and *in vitro* progenitor capacity.

In the absence of established procedures to enrich human Club/Hillock cells from prostate specimens, the progenitor-like properties of the latter cells have never been experimentally assessed. We here identified the HPV-10 cell line as the best surrogate of tumoral Club/Hillock cells. According to their prostatic adenocarcinoma origin (Hukku *et al*., 2000) and their enrichment in the LSC^med^-0 signature, HPV-10 cell cells appear to be a relevant model of Club-like cells preexisting in naïve tumors. We show that these cells are able to form tumorspheres *in vitro,* which is consistent with the *in silico*-predicted progenitor-like capacities of Club/Hillock cells (Henry *et al*., 2018). As observed with *Pten*^pc-/-^ LSC^med^ cells, combined JQ-1/CX-6258 treatment completely abrogated the tumorsphere-forming capacity of HPV-10 cells. Their growth was also markedly reduced while their viability was mildly affected. Taken together, the results of our *in vitro* and *in vivo* assays suggest that combined pharmacological inhibition of the FOSL1/AP-1 complex and PIM1 interferes with the growth, survival, transcriptomic reprogramming and progenitor capacities of mouse and human Club-like cells. Further studies are needed to understand how all these effects work together to abrogate the promotion of CRPC growth by Club-type cells.

Neuroendocrine differentiation (NED) is a mechanism of late-stage therapeutic resistance, increasingly observed since the generalization of ARPI treatment of advanced prostate cancer (Yamada & Beltran, 2021). Several observations suggest that LSC^med^/Club/Hillock cells may be predisposed to NED under therapeutic pressure. *Wfdc2* and *Sox2* are part of the LSC^med^ cell signature (Baures *et al*., 2022a); both genes are positively associated with neuroendocrine signature in human PCa (Dong *et al*, 2020), and *Sox2* is essential for NED in mice (Kwon *et al*, 2021). Similarly, human Club/Hillock cells are enriched in neuronal stem programs (Yan *et al*, 2022). In *Pten*^pc-/-^ mice, our scRNAseq analysis showed that LSC^med^-2 cells, which are moderately amplified after castration, are characterized by the expression of *Onecut2*. This transcription factor is a major inducer of early NED and biochemical recurrence via the inhibition of AR and FoxA1 transcriptional programs (Rotinen *et al*., 2018). Still, LSC^med^-2 cells showed only low enrichment in NEPC signature and no expression of late NED markers 2 months post-castration. This is consistent with the unaltered LSC^med^-2 signature observed by spatial transcriptomics analysis. The heterogeneity of LSC^med^-2 marker appearance after castration (RT-qPCR data) is informative as it suggests that the kinetics of progression towards the LSC^med^-2 state is not only delayed, but also animal-dependent, compared to the emergence of the LSC^med^-1 state. Together, these data suggest that a longer delay than the 2 months post-castration analyzed in this study may be required to alter the expression level of factors known to positively regulate NED (e.g. *Ascl1* or *Ezh2*) or negatively (*Klf5*). Further genetic stress, e.g. loss of *Rb1* and/or *Tp53,* two key negative regulators of the neuroendocrine signature (Qian *et al*, 2022), may also be required to drive full NED of some LSC^med^-2 cells. Of interest, Onecut2 was recently shown to be a broadly acting lineage plasticity facilitator as it also supports the appearance of treatment-resistant adenocarcinoma through multiple mechanisms (Qian *et al*., 2024). In this context, strategies targeting progenitor cells presenting with the LSC^med^-2 profile may also be relevant to prevent progression towards aggressive forms of CRPC. The consistent expression of AP-1 family members and PIM kinases in LSC^med^-2 cells suggests that our targeting strategy using JQ-1/CX-6258, primarily designed to target LSC^med^-1 cells, may also counteract LSC^med^-2 cell amplification post-castration. Alternatively, the recently reported small molecule inhibitor of ONECUT2 (Qian *et al*., 2024) should also be considered. Additional studies are needed to elucidate the actual kinetics of LSC^med^-2 cell emergence in *Pten^pc-/-^* mice in order to challenge these therapeutic strategies.

This study presents some limitations. While *FOSL1* silencing *in vitro* confirmed the relevance of targeting this transcription factor in Club-like cells, additional genetically-engineered models, including *Fosl1*-deficient *Pten^pc-/-^* mice, are needed to better delineate the actual effects of FOSL1 inhibition on LSC^med^ cell growth, survival, plasticity and progenitor properties. Such models will also help identify any functional redundancy involving other AP-1 members, which may promote therapeutic resistance to FOSL1 inhibition. Another hurdle relates to the specificity of the drugs used to target FOSL1. Although JQ-1 was shown to mainly target FOSL1 and to phenocopy *FOSL1* knockdown in various tumors (Baker *et al*., 2015; Bid *et al*., 2016; Casalino *et al*., 2022; Lockwood *et al*., 2012), other genes may also be altered by this BET/AP-1 inhibitor. In this matter, exhaustive *in vitro* and *in vivo* characterization of the effects of T5224-PROTAC, recently shown to directly bind to, and degrade, FOSL1 (Zaman *et al*., 2024), is necessary to evaluate its therapeutic superiority to JQ-1 in terms of specificity and efficacy.

In conclusion, our study uncovers the critical role of transcriptional reprogramming in pre-existing Club-like cells as a novel mechanism driving castration resistance. We demonstrate that FOSL1/AP-1 is a key regulator of this molecular transformation, resulting in the emergence of aggressive progenitor cells enriched in basal characteristics. Furthermore, our findings align with recent research indicating that the shift from androgen-dependent luminal cells to androgen-independent progenitor-like cells during castration is similarly mediated by this transcriptional complex. This establishes FOSL1/AP-1 as a pivotal factor in the mechanisms underlying resistance to anti-androgen therapy. Therefore, our dual therapeutic approach targeting both FOSL1/AP-1 and PIM kinases combined to castration presents a promising strategy to counteract these early mechanisms of castration resistance and disrupt the progression towards CRPC.

## Methods

### Reagents and Tools Table

See enclosed file

### Methods and Protocols

#### Animals

Mouse colonies were housed in controlled conditions, on a 12/12-h light/dark cycle with normal food and water provided ad libitum. *Pten^pc-/-^*male mice were generated by breeding *Pten*^loxP/loxP^ female mice with Pb-Cre4/*Pten*^loxP/loxP^ transgenic males and maintained on a mixed C57BL/6 and Sv/129 genetic background as described previously (Baures *et al*., 2022b; Sackmann Sala *et al*., 2017). Experiments were performed using 7- to 11-month-old mice, i.e. when aggressive malignant phenotypes were well established. As indicated, mice were surgically castrated and prostate were analyzed between 5 days and 2 months following castration. Prostate samples were obtained by microdissection immediately after sacrifice by cervical dislocation. Under a dissection microscope, adipose tissues were removed from the urogenital tract. The bladder, the ampullary gland and the urethra were removed to isolate the four prostate lobes. All animal procedures have been extensively described in previous publications (Baures *et al*., 2022b; Sackmann Sala *et al*., 2017).

Breeding and maintenance of mice were carried out in the accredited animal facility of the Necker campus in compliance with French and European Union regulations on the use of laboratory animals for research. Animal experiments were approved by the local ethical committee for animal experimentation (APAFIS authorizations #40276 and #49662).

#### JQ-1 and CX-6258 Treatments

JQ-1 (Cat# HY-13030) and CX-6258 (Cat# HY-18095) were purchased from MedChemExpress. For *in vivo* treatments, 7-month-old *Pten^pc−/−^*mice were castrated, then after 4-day recovery, they were randomized in two groups (n= 8 mice each) to receive either vehicle or a combination of JQ-1 (daily intraperitoneal injection, 50mg/kg solubilized in 5% DMSO, 40% PEG300, 5% Tween80 and 50% H_2_O) and CX-6258 (biweekly oral administration, 100mg/kg solubilized in 15% Cremophor and PBS), for 4 weeks. After euthanasia, the prostates were immediately microdissected and weighed, then fixed or processed for cell sorting, protein or RNA extraction, as described below.

For human PCa cell xenograft experiments, we used 12-week-old NSG male mice (NOD.Cg-Prkdc^scid^ Il2rg^tm1Wjl^/SzJ) purchased from Charles River Laboratories International Inc.. Animals were castrated and injected subcutaneously with 1.10^6^ PC-3 (ATCC, cat# CRL-1435, RRID: CVCL_0035) in Matrigel on both flanks to fulfil the 3R rule. Four days later, mice were randomized in two groups of 6 mice each (n=12 tumors) to receive vehicle or combined CX-6258 and JQ-1 treatment as described above. Before tumors were measurable, two mice in the treated group experienced issues with the gavage procedure that prevented continuation of oral treatment for ethical reasons. From day 15 of treatment, tumors were measured twice a week using a digital caliper. Tumor size was calculated using the formula (a × b2)/2, where a = length and b = width of the tumor. One tumor in the treated group initially grew abnormally fast (outlier) and regressed at the last timepoint (Appendix Fig. S6), therefore it was not included in the analysis. After euthanasia, tumors were dissected, weighed and photographed. Tumors were then embedded in paraffin for further analysis. Investigators who performed mouse analyses were blinded to mouse treatments (vehicle *versus* drugs).

#### Histological Classification Post-Treatment and Treatment Toxicity

To evaluate the effect of treatments on prostate histology, prostates from *Pten^pc−/−^* mice were fixed in 4% PFA, paraffin wax-embedded, and sections were stained with hematoxylin and eosin (H&E). Analysis was performed by using the QuPath Software. Glands and lumen were manually detected and outlined. For each gland, we measured the number of lumens and the proportion of lumen to the total surface of the gland. Based on these parameters, glands were classified in 4 different categories: (1) normal gland (a single lumen representing >70% of the gland), (2) PIN structure (a single lumen representing <70% of the gland), (3) cribriform structure (two or more lumens representing >80% of the gland) and (4) malignant gland (two or more lumens representing <80% of the gland or absence of any lumen). Following this analysis, normal and PIN structures were classified as “benign” (*versus* “tumoral”). Investigators who performed mouse analyses were blinded to mouse treatments (vehicle *versus* drugs).

To evaluate the potential toxic effects of the administered drug combination, we analyzed organ samples stained with H&E. The kidneys, lungs, and livers from five *Pten^pc-/-^* mice randomly-chosen in both the control and treated groups were examined. Two pathologists blinded to group assignments independently performed the histological analysis. Degenerative and inflammatory lesions were rated on a 5-point scale (0 = no lesion, 1 = scattered, 2 = mild, 3 = moderate, 4 = marked). These analyses were complemented by quantification of cell death using the TUNEL assay described below.

#### Spatial Transcriptomics

Digital Spatial Profiling (GeoMx™, Bruker) was performed on prostate sections from two naïve *Pten^pc-/-^* mice, two castrated and vehicle-treated *Pten^pc-/-^* mice, and two castrated and JQ-1 + CX-6258-treated *Pten^pc-/-^* mice. Five µm sections were baked for 30 min at 60 °C. The nuclei and the epithelial cells were stained using SYTO™ 83 Orange Fluorescent Nucleic Acid Stain (S11364, Thermo Fischer) and the Alexa Fluor 488 anti-mouse CD236 (EpCAM) antibody (#AB237384, dilution 1/100, Abcam), respectively. *In situ* hybridization of RNA-directed DNA oligo probes (Nanostring Mouse Whole Transcriptome Atlas) was performed according to the manufacturer’s protocol. For each mouse, the EpCAM+ (i.e. epithelial) cells in 5 to 7 regions of interest (ROI) in the DLP and 5 in the AP were selected. The signature score was calculated using the normalized count obtained from the GeoMx pipeline and the AddModuleScore function from Seurat. The deregulated genes were considered for a p-value < 0.05 and a log2 fold change of 0.5.

#### Prostate dissociation

Prostates were minced using razor blades and digested in a solution of Dulbecco’s modified eagle medium (DMEM) (Thermo Fisher Scientific) containing 10% FBS (Eurobio), 1% penicillin/streptomycin (Pen/Strep) (Thermo Fisher Scientific) and 1 mg/mL Collagenase Type I solution (Thermo Fisher Scientific) for 1h at 37°C, followed by 5-minute incubation at 37°C in 0.25% Trypsin (Thermo Fisher Scientific). The digestion was stopped with a solution of DMEM containing DNase I (Roche). Cells were passed 10 times through a 20G syringe then filtered through a 40 µm cell strainer to generate a single-cell suspension. Cells were subjected to differential centrifugation using Histopaque-1119 (Sigma-Aldrich) to reduce the overall level of secretion in the sample. An aliquot of cells was stained with Trypan blue (Thermo Fisher Scientific) and counted using a hemocytometer to assess the cell viability.

#### FACS-Sorting of LSC^med^ Cells

The procedure for cell sorting was performed as previously described (Lukacs *et al*, 2010; Sackmann-Sala *et al*, 2014; Sackmann Sala *et al*., 2017). Isolated prostatic cells were stained for FACS on ice for 30 minutes using the following rat antibodies (all from eBioscience): fluorescein isothiocyanate-coupled lineage (Lin) antibodies (anti-CD31, -CD45 and -TER-119; dilution 1/500), phosphatidylethanolamine-Cyanine7-coupled anti-EpCAM,(1/500), phosphatidylethanolamine-coupled anti-CD49f (integrin alpha-6; dilution 1/50) and allophycocyanin-coupled anti-Sca1 (lymphocyte antigen 6A-2/6E-1; dilution 1/150). Dead cells were stained using SYTOX Blue (Life Technologies). Cell sorting was performed on a BD FACS Aria III (BD Biosciences) in DMEM containing 2% FBS and 1% Pen/Strep. Lineage antibodies (CD31, CD45, TER-119) were used to deplete hematopoietic, endothelial and immune cells and EpCAM antibody to distinguish epithelial from stromal cells. CD49f and SCA-1 markers were used to select LSC^med^ cells from the other epithelial cell types. Sorted cells were collected in DMEM medium supplemented with 50% FBS and 1% Pen/Strep.

#### Single-cell RNA-Sequencing and Analyses

LSC^med^ cancer cells sorted from 2 pooled intact and 2 pooled castrated *Pten^pc-/-^* mice were dispensed (BD FACS Aria III) in 96 well plates (VWR, DNase, RNase free) containing 2 μL of lysis buffer (0.2% Triton X-100, 4U of RNase inhibitor, Takara) per well. Plates were properly sealed and spun down at 2,000 g for 1 min before storing at −80°C. Whole transcriptome amplification was performed with a modified SMART-seq2 protocol as described previously (Picelli *et al*, 2014) using 23 instead of 18 cycles of cDNA amplification. PCR purification was realized with a 0.8:1 ratio (ampureXP beads:DNA). Amplified cDNA quality was monitored with a high sensitivity DNA chip (Agilent) using the Bioanalyzer (Agilent). Sequencing libraries were performed using the Nextera XT kit (Illumina) as described previously (Picelli *et al*., 2014) using 1/4^th^ of the recommended reagent volumes and 1/5^th^ of input DNA with a tagmentation time of 9 min. Library quality was monitored with a high-sensitivity DNA chip (Agilent) using the Bioanalyzer (Agilent). Indexing was performed with the Nextera XT index Kit V2 (A-D). Up to 4 × 96 single cells were pooled per sequencing lane. Samples were sequenced on the Illumina NextSeq 500 platform using 75bp single-end reads.

#### Single-Cell Transcriptome Analysis

The merged gene expression matrix (raw counts) containing all samples was analyzed with the Seurat3 package and Seurat_Extend (Hua *et al*, 2025). Cells were filtered by nFeature_RNA (genes detected) >3000 and <7000, percent.mt (percentage of mitochondria genes) < 10. This resulted in n=306 valid cells for downstream analysis. After filtering cells, log-normalization was performed using the default NormalizeData function in Seurat. The scaled data was regressed for cell cycle phases (S.Score and G2M.Score) and number of genes per cell (nFeature_RNA) before performing UMAP based dimension reduction (dims=1:20, resolution =0.4).

#### StemChecker Analysis

Seurat cluster (LSC^med^-0, LSC^med^-1, LSC^med^-2) characteristic murine genes (adj.pval>0.05) were analyzed for “stemness” enrichment using the stemchecker tool (RRID:SCR_025014), while masking cell cycle genes.

#### Pseudotime-Ordering Analysis

The raw count matrix (n=306 cells) was used as input for Monocle analysis (2.14.0) (RRID:SCR_016339) and normalized by the M3Drop function. Dimension reduction was based on the variable genes (n=1483) (max_components = 2, method = ‘DDRTree’). Ordering of the cell was performed by the f orderCells (HSMM_myo, num_paths = 2, reverse = T) function and the resulting trajectory was colored by pseudotime, cell type and Seurat cluster information. Differential gene expression analysis between groups through a likelihood ratio test for comparing a full generalized linear model with additional effects to a reduced generalized linear model based on negative binomial distributions.

#### Differentiation Trajectory Analysis Using Velocyto and scVelo

To infer the directionality of differentiation, we employed the Velocyto pipeline (La Manno *et al*, 2018). Utilizing the *.bam files and cell metadata, we computed the spliced and unspliced RNA matrices. Differentiation trajectories were visualized using UMAP embeddings, which were generated with Seurat in R (RRID:SCR_016341). The trajectories, represented by arrows, were plotted using ScVelo (Bergen *et al*, 2020) in Python.

#### Gene set enrichment analyses

Gene enrichment analyses were conducted for each Seurat cluster by interrogating the C2 Chemical and Genetic Perturbations category of MsigDB via the hypeR (2.2.0) package and the GeneSetAnalysis function from the SeuratExtend package.

We employed the Hallmark 50 gene set, a collection of well-defined, biologically relevant gene sets from the Molecular Signatures Database (MSigDB, https://www.gsea-msigdb.org/gsea/msigdb/human/genesets.jsp?collection=H). This analysis was performed to identify significantly enriched pathways across the LSC^med^-0, LSC^med^-1, and LSC^med^-2 clusters. The results were visualized using a heatmap, with color intensity representing z-score normalized enrichment scores.

#### SCENIC analysis

To investigate regulon networks, we employed a comprehensive approach combining computational analysis and visualization techniques. We utilized the SCENIC algorithm (https://github.com/aertslab/SCENIC) to infer gene regulatory networks and regulon activities. The analysis pipeline was implemented using the SeuratExtend package (Hua *et al*, 2024) for data processing and visualization. We generated waterfall plots to highlight differential TF regulon activities between selected clusters and other cells, with the top 10 TFs labeled for clarity. Gene regulatory networks were visualized in Cytoscape (http://cytoscape.org), with nodes representing genes (round nodes) and TFs (square nodes). Node colors were mapped to relative gene expression or regulon activity using z-score normalization, enabling comparison of expression and activity patterns among LSCmed-0, LSCmed-1, and LSCmed-2 clusters. To manage the visualization of large regulons, we capped the number of target genes per transcription factor (TF) at 200, prioritizing the most variable genes based on their ranking from Seurat’s FindVariableFeatures function. This approach ensured a balance between capturing key regulatory relationships and maintaining analytical tractability.

#### Cell Lines and Treatments

HPV-10 (ATCC, Cat# CRL-2220), PC-3 (ATCC, cat# CRL-1435,) and LNCaP (ATCC, cat# CRL-1740) PCa cell lines were purchased from ATCC (Manassas, USA) at the beginning of this study. Frozen stocks were generated between passages 2 to 5 of initial cell cultures, then aliquots were thawed for each experiment and used before passage 20. Therefore, no cell authentication was performed. None of the cells used in this study are listed in the ICLAC database of commonly misidentified lines (https://iclac.org/databases/cross-contaminations/).

Cells were grown at 37°C and 5% CO_2_. HPV-10 cells were cultured in Keratinocyte Serum Free Medium (K-SFM) (Thermo Fisher Scientific, Cat# 17005-034) supplemented with 0.05mg/mL Bovine Pituitary Extract (BPE) (Thermo Fisher Scientific, Cat# 13028-014), 5ng/mL EGF (PeproTech, Cat# AF-100-15-500UG) and 1% Pen/Strep. PC-3 and LNCaP cells were cultured in DMEM (Thermo Fisher Scientific, Cat# 31966-021) supplemented with 10% FBS (Eurobio Scientific, Cat# CVFSVF00-01, #lot S80515) and 1% Pen/Strep. Medium was changed every other day.

For drug treatments, HPV-10 and PC-3 cells were plated in 6-well plates at 50,000 and 30,000 cells/well, respectively. After 3 days, cells were treated with DMSO, JQ-1 and CX-6258, alone or combined, and analyzed after 72h, according to the protocols described below. Alternatively, HPV-10 and PC-3 cells were plated in 24-well plates at 50,000 cells/well. The next day, cells were treated with DMSO or T5224-PROTAC (5 or 15µM) at 0, 8 and 24h, and analyzed at 48h.

For siRNA transfections, PC-3 cells were plated in 24-well plates at 30,000 cells/well. The next day, cells were treated with Lipofectamine (Thermo Fisher Scientific, Cat# 18324-012) with siScrambled or siFOSL1 in OPTIMEM (Thermo Fisher Scientific, Cat# 31985-070) for 6h before changing medium back to complete medium. Cells were analyzed at 72h.

For cell number analysis, wells were rinsed in PBS and adherent cells were trypsinized before being manually counted using a Malassez counting cell. For cell viability analysis, culture medium (containing floating cells) and adherent cells (trypsinized) were collected, centrifuged (300g) and resuspended in HBSS before being counted with trypan blue as above. Cell viability was quantified as the ratio of unstained *versus* total (unstained + stained) cells.

#### Reverse Transcription-Quantitative PCR (RT-qPCR)

RNA extractions from *Pten^pc−/−^* LSC^med^ cells and human PCa HPV-10 cells were performed with the Nucleospin RNA XS and Nucleospin RNA (Macherey-Nagel), respectively, following the manufacturer’s protocols. Reverse transcription was carried out using the SuperScript™ VILO™ cDNA Synthesis Kit (Invitrogen) for murine cells and the GoScript™ Reverse Transcriptase (Promega) for the human cell line, according to the manufacturer’s instructions.

For qPCR, iTaq Universal SYBR Green Supermix (Promega) was used, and reactions were run on a qTower 2.0 real-time thermal cycler (Analytik Jena). Expression data were normalized to Cyclophilin A for murine samples and Actin B for human samples.

Primer sequences (Sigma-Aldrich) are listed in Appendix Table S3.

#### Immunoblotting

Cell lysates were prepared on ice in an appropriate lysis buffer (50 mM Tris-HCl, pH 7.5, 150 mM NaCl, 1% TRITON X-100, 0.5% NP-40, 10% glycerol, 1% protease, and a Phosphatase Inhibitor Cocktail (78442, Thermo Fisher Scientific)). Protein concentrations were determined with the Pierce BCA protein assay kit (23225, Thermo Fisher Scientific). Equal protein amounts (30–40 µg) diluted in a 4× Laemmli buffer were denatured by heating at 95 °C for 5 min and separated by electrophoresis on 4–12% NuPAGE Bis-Tris Gel, and then transferred onto a 0.45μm PVDF membrane. Membranes were blocked with 5% non-fat dry milk in PBS-T (PBS with 0.1% Tween-20) for 1 h at room temperature and then incubated with primary antibodies at 4 °C overnight. Membranes were then washed with PBS-T, and incubated with the appropriate HRP-coupled secondary antibody for 1 h 30 min at RT. Membranes were then washed with PBS-T, and bound antibodies were detected using an ECL detection kit (Immobilon Western ECL, Millipore) and ChemiDoc Imaging Systems (BioRad) using the CCD camera for light capture according to the manufacturer’s protocol. Signals were quantified using Image Lab Software (Bio-Rad) and normalized to GAPDH.

#### Organoid and tumorsphere culture and treatment

We used the reference protocol for prostate organoid culture described by Clevers’ lab (Drost *et al*, 2016) with modifications. LSC^med^ cells sorted from *Pten*^pc−/−^ mice and human PCa HPV-10 cells were plated in triplicate on a Low Growth Factor-containing Matrigel (Corning) layer in a 96-well plate (Sarstedt) at concentrations of 3,000 or 1,000 cells/well, respectively, and were cultured at 37°C and 5% CO_2_. After one day of incubation, the culture medium was removed and the cells were covered by a new layer of Matrigel to generate 3D cell structures. The organoid-forming capacity did not differ from the efficacy obtained using 3D droplet culture earlier described (Drost *et al*., 2016). Drugs were added at this stage and the medium (+ drugs) was changed every other day. After 10 days of culture, organoids were fixed in 4% PFA and images were taken with a 4x objective on an M5000 EVOS inverted microscope (Invitrogen) to capture the entire surface of the well. Organoid counting and surface area measurements were performed using Fiji software (http://fiji.sc) by manually outlining the organoid surface. Only structures with size superior to 2000μm^2^ were considered as organoid. The number and size of organoids in the various experimental conditions were normalized to cognate mean value obtained in the non-treated condition.

#### Immunohistochemistry (IHC) and Immunofluorescence (IF)

For IHC, all murine prostate samples were fixed in 4% PFA, paraffin wax-embedded, and sections underwent heat-induced antigen retrieval in citrate buffer at pH 6 (95°C, 30 min). IHC was performed as previously described (Baures *et al*., 2022b) using antibodies directed against KI-67 (Zytomed Systems, Cat# RBK027-05; dilution 1/100), FOSL1 (Santa-Cruz Biotechnologies, cat# sc-376148, dilution 1/10) and CD44 (Biolegend, Cat#, cat#103001, dilution 1/100). Signal was amplified and detected using the Vector Elite ABC HRP kit with DAB substrate (Vector Laboratories) and nuclei were counterstained with hematoxylin. Slides were scanned with a Nanozoomer 2.0 (Hamamatsu) and analyzed using NDP.view 2 software (Hamamatsu). Positive nuclei were quantified in 10 fields per slide using QuPath Software (https://qupath.github.io/).

For IF, cells were fixed in PFA 4% for 15 minutes at room temperature and permeabilized for 15 minutes in PBS supplemented with 0.2% Tween-20. Rabbit anti-KRT4 (Abcam, ab51-599, dilution 1/100) and anti-KRT13 (Sigma-Aldrich, HPA030877, dilution 1/200) antibodies were incubated overnight at 4°C and secondary antibodies were incubated during 1h at room temperature. Nuclei were stained with Hoechst dye. Samples were analyzed using a 40× objective under an Apotome 2 microscope (Zeiss).

#### Terminal Deoxynucleotidyl Transferase dUTP Nick End (TUNEL) Labeling

The TUNEL assay was performed with the *In Situ* Cell Death Detection Kit (Roche, Cat. #11 684 809 910) according to the manufacturer’s instructions. Briefly, samples underwent dewaxation and rehydratation. Slides were heated in Citrate Buffer during 5 minutes at 350W and then incubated for 1h at 37°C, with the TUNEL reaction mixture. Fluorescence was observed and images were taken using a 20x objective under an M5000 EVOS inverted microscope.

#### Statistical analyses

Unless otherwise stated, the data describe biological replicates, and the number of independent experiments is indicated in Figure captions. Error bars represent S.D. The statistical tests performed are indicated in figure captions for each experiment (ns, not significant; *p<0.05; **p<0.01; ***p<0.001; ****p<0.0001). A value of p < 0.05 was used as significance cutoff for all tests. Statistical analyses were performed using GraphPad Prism version 9.00 for Windows (http://www.graphpad.com/). Statistical parameters used for transcriptomic data analyses are described in the corresponding Methods section. Animals were randomized in experimental groups. Investigators who performed histopathological analyses of mouse tissues were blinded to mouse treatments (vehicle *versus* drugs).

## Data and materials availability

All data needed to evaluate the conclusions of this study are presented in the main text and the Supplementary Materials. Computational analyses were based on already published pipelines. The detailed code is available upon request. The datasets produced in this study are available in Gene Expression Omnibus repository:

- scRNAseq of *Pten^pc-/-^* LSCmed cells: GSE273079 (private status: ovozmciyhtonjux) https://www.ncbi.nlm.nih.gov/geo/query/acc.cgi?acc=GSE273079
- Digital Spatial Profiling of *Pten^pc-/-^* prostate sections: GSE302983 (private status: kxgneqayxjezvgz) https://www.ncbi.nlm.nih.gov/geo/query/acc.cgi?acc=GSE302983

## Acknowledgments

The authors are grateful to the personnel of the technological core facilities of the SFR Necker including animal housing (in particular Emilie Panafieu, Jean-Chistophe Beche, Razack Alao, Emeline Giton and Clément Seigneurin, Auguste Ellong Mbongo), cytometry (in particular Jérôme Mégret), histology, and image analysis (Nicolas Goudin). They warmly thank Sandra Högler for help in histopathological assessments. We thank Romain Kaiser, Tao Ye and Christelle Thibault-Carpentier from GenomEast, a member of the “France Génomique” consortium (ANR-10-INBS-0009) for the technical support for the digital spatial profiling.

This research was funded by Institut National du Cancer, grant INCa_16077 (VG & DM); Ligue contre le cancer, grants RS18/75-48, RS19/75-63, RS20/75-93 and RS21 /75-35 (VG); Association pour la recherche sur les tumeurs de la prostate (VG); FONCER contre le cancer (JEG); Annual funds from Inserm and CNRS (VG & DM); University Paris Cité (VG); Interdisciplinary Thematic Institute IMCBio+, as part of the ITI 2021-2028 program of the University of Strasbourg (DM); IdEx Unistra (ANR-10-IDEX-0002) (DM); SFRI-STRAT’US project (ANR-20-SFRI-0012) (DM); EUR IMCBio (ANR-17-EURE-0023) under the framework of the France 2030 Program (DM).

M.B. and A-S. V.A. were supported by a fellowship from the Ministry of Research, M.B. and A. K. by INCa_16077 grant, C.D. by a PhD fellowship Aviesan (Alliance nationale pour les sciences de la vie et de la santé)/ITMO Cancer supported by Inserm, and V.F. by a PhD fellowship from Inserm (Ecole de l’Inserm Pfizer Innovation). F.R. is funded by the Melanoma Research Alliance and the Wolfgang & Gertrud Boettcher Foundation.

## The paper explained

### Problem

The major challenge in prostate cancer treatment is to overcome resistance to castration therapy that ultimately leads to patient death. The mechanisms that drive the shift of tumors from castration-sensitivity to castration-resistance remain poorly understood. Club cells are a newly-identified progenitor-like cell type predicted to be castration-resistant. While these cells have been identified in prostate tumors prior to castration, their functional contribution to therapy resistance is unexplored.

### Result

In this study, we used a mouse model of castration-resistant prostate cancer (prostate-specific inactivation of the tumor suppressor gene *Pten*) that is enriched in Club-like cells. We report that the castration tolerance of these cells is not intrinsic, but results from a transcriptional shift that enhances several hallmarks of cancer aggressiveness such as epithelial-to-mesenchymal transition, basal cell features and stem-like programs. We identified the transcription factor FOSL1/AP-1 as a major driver of this process, called cell plasticity. Dual pharmacological targeting of FOSL1 and PIM1, a serine/threonine kinase known to promote prostate cancer, abrogates the *in vitro* progenitor properties of the various mouse and human Club-like models that we used, and this was correlated to a marked reduction of *in vivo* tumor growth after castration.

### Impact

Our findings show that targeting stemness and tumor-initiating capacities of Club-like prostatic cells by dual inhibition of FOSL1/AP-1 and PIM kinase is a promising strategy to potentiate androgen deprivation therapy and mitigate prostate cancer relapse.

## Expanded View (EV) Figures

**Figure EV1.**
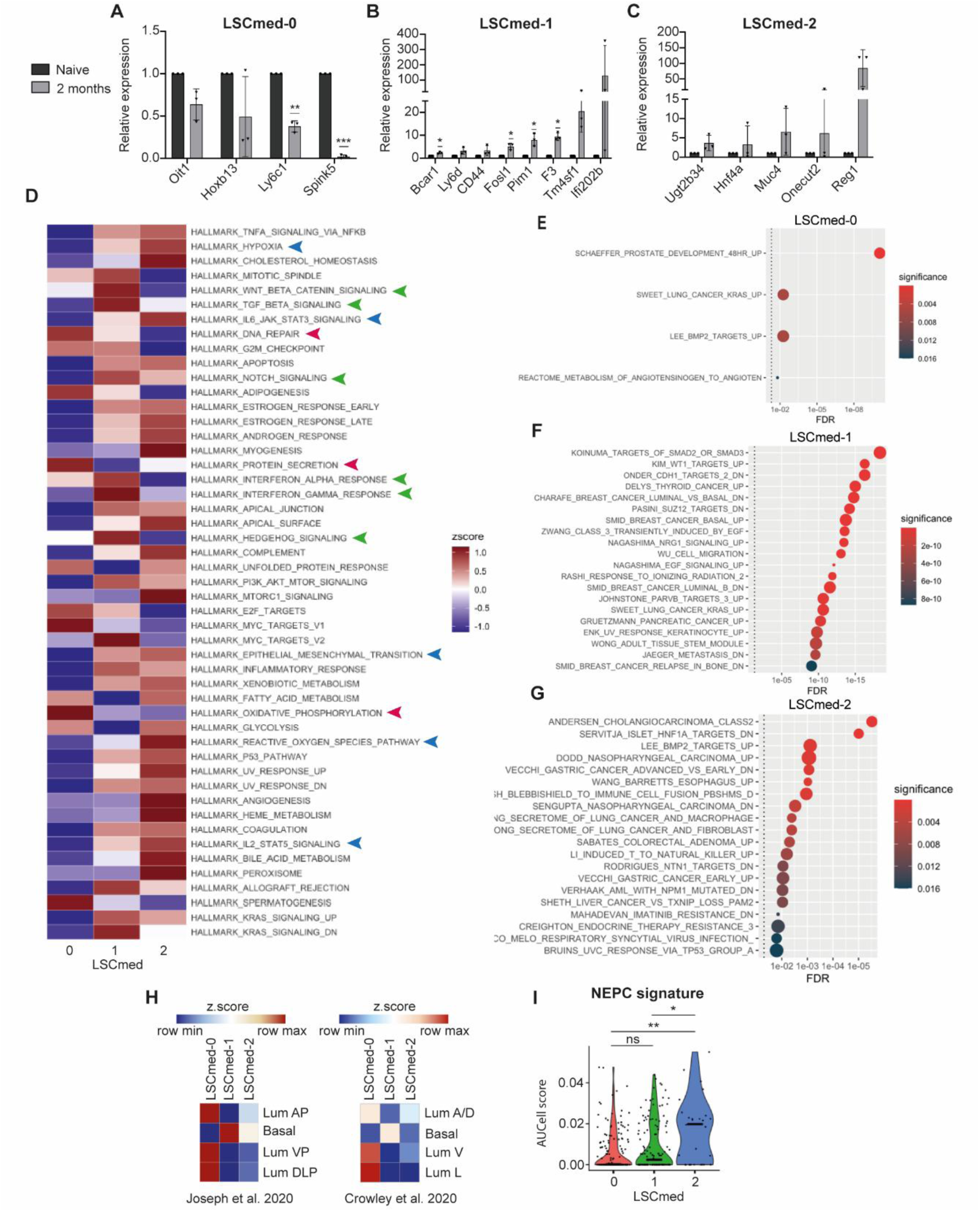
Gene set enrichment analyses of the three LSC^med^ cell subpopulations. **(A-C)** The expression of selected marker genes of LSC^med^-0 **(A)**, LSC^med^-1 **(B)** and LSC^med^-2 **(C)** cells was measured by RT-qPCR in bulk LSC^med^ cells sorted from intact (n=3) and 2-month-castrated (n=7) Pten^pc-/-^ mice (biological replicates, n = 3 independent experiments). Data are normalized to the expression in intact mice. ns, not significant, *p<0.05 (unpaired t test with Welch’s correction). Exact p-values are reported in Appendix Table S1. **(D)** Heatmap highlighting Hallmark 50 gene set enrichments of the three LSC^med^ subpopulations. Biological processes mentioned in the text are labelled with arrows color-coded according to Figure 2A (red: LSC^med^-0; green LSC^med^-1; blue LSC^med^-2). **(E-G)** Dot plots showing enrichment of gene sets belonging to the C2 Chemical and Genetic Perturbations category (MsigDB) per LSC^med^ cluster using the hypeR (2.2.0) package. **(H)** Heatmap representation of LSC^med^-0, LSC^med^-1 and LSC^med^-2 signatures (average expression, z.score) in luminal and basal cells. We interrogated murine prostate scRNA-seq data sets (Data Ref: Crowley *et al*., 2020a; Joseph *et al*., 2020a) for the expression of top50 upregulated LSC^med^ cluster 0,1,2 genes. The average LSC^med^ signature expression was determined per cell type and z.score-transformed. LSC^med^ cluster 0,1,2 z.scores were plotted for luminal (Lum) and basal cell types. Abbreviations: A (anterior), D (dorsal), L (lateral) and V (ventral) prostate lobes. **(I)** NEPC signature expression intensities (AUCell score) in LSC^med^ cell subpopulations. ns, not significant, *p<0.05, **p<0.01 (Wilcoxon test, Holm-method adjusted). Exact p-values: p=0,12 (0 vs 1), p=0,0031 (0 vs 2) and p=0,039 (1 vs 2).

**Figure EV2.**
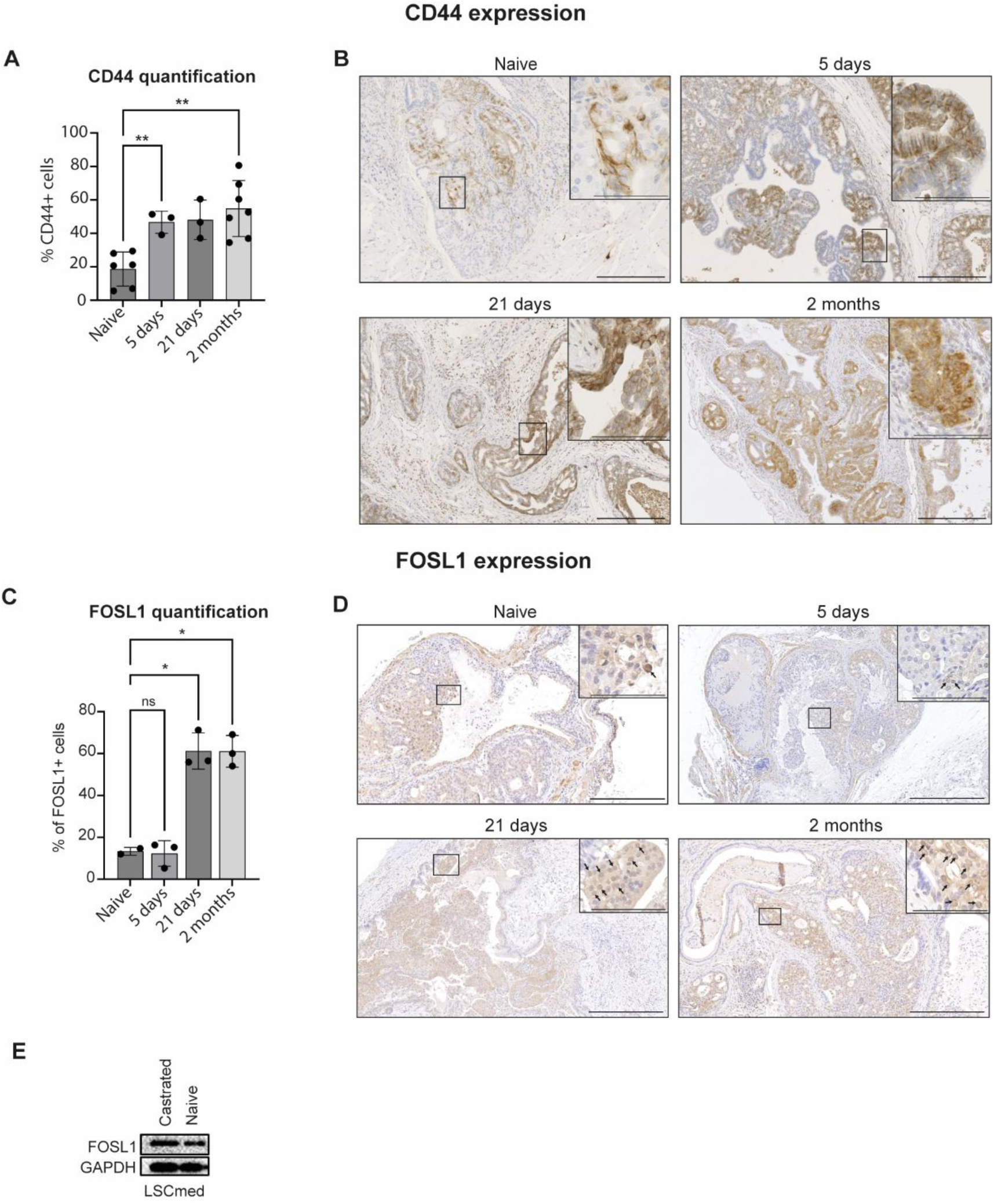
Time-course analysis of CD44 and FOSL1 expression after castration in *Pten^pc-/-^*mouse prostates. **(A,B)** The expression of CD44 chosen as a selected marker of LSC^med^-1 cells was assessed by immunohistochemistry in intact *Pten^pc-/-^* mice (n=6) and 5 days (n=3), 21 days (n=3) and 2 months (n=7) post-castration. Both membrane and diffuse cytoplasmic staining was observed as previously reported in PCa (Omara-Opyene *et al*, 2004). **(A)** Quantification of CD44-positive cells was performed by QuPath. *p<0.05, **p<0.01 (Welch’s ANOVA with Brown-Forsythe correction, Dunnett’s T3 post hoc test). Exact p-values: p=0.0071 (5 days), p=0.0523 (21 days), p=0.0022 (2 months). **(B)** Representative images are shown for each time point. Scale bars = 200 µm and 50 µm in insets. **(C,D)** The expression of FOSL1 was assessed by immunohistochemistry in intact *Pten^pc-/-^* mice (n=2) and 5 days (n=3), 21 days (n=3) and 2 months (n=3) post-castration. Both nuclear and diffuse cytoplasmic staining was observed as previously reported in cancer cells (Sobolev *et al*., 2022; Song *et al*., 2006; Taha *et al*., 2023). Arrows point to some positive nuclei. **(C)** Quantification of FOSL1-positive cells (nuclear + cytoplasmic) was performed by QuPath. ns, not significant, *p<0.05 (Welch’s ANOVA with Brown-Forsythe correction, Dunnett’s T3 post hoc test). Exact p-values: p=0.9883 (5 days), p=0.0239 (21 days), p=0.0190 (2 months). **(D)** Representative images are shown for each time point. Scale bars = 200 µm and 50 µm in insets. **(E)** Western blot showing upregulation of FOSL1 expression in sorted LSC^med^ cells after castration (+46%).

**Figure EV3.**
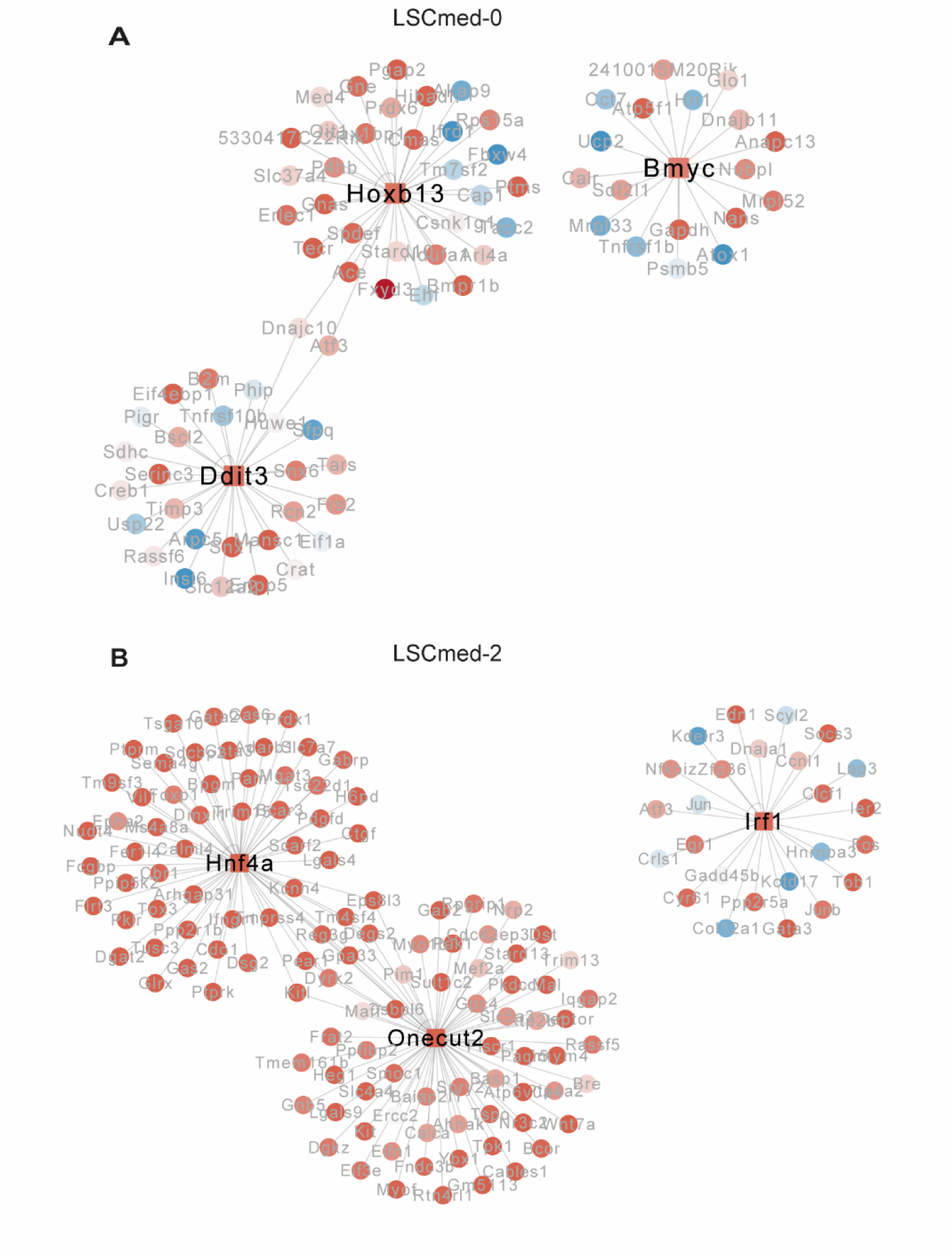
Transcription factor networks in LSC^med^-0 and LSC^med^-2 cells. The transcription factor networks of the 3 most pertinent regulons of LSC^med^-0 **(A)** and LSC^med^-2 cells **(B)** are represented. See Fig. 4C showing the same analysis for LSC^med^-1 cells.

**Figure EV4.**
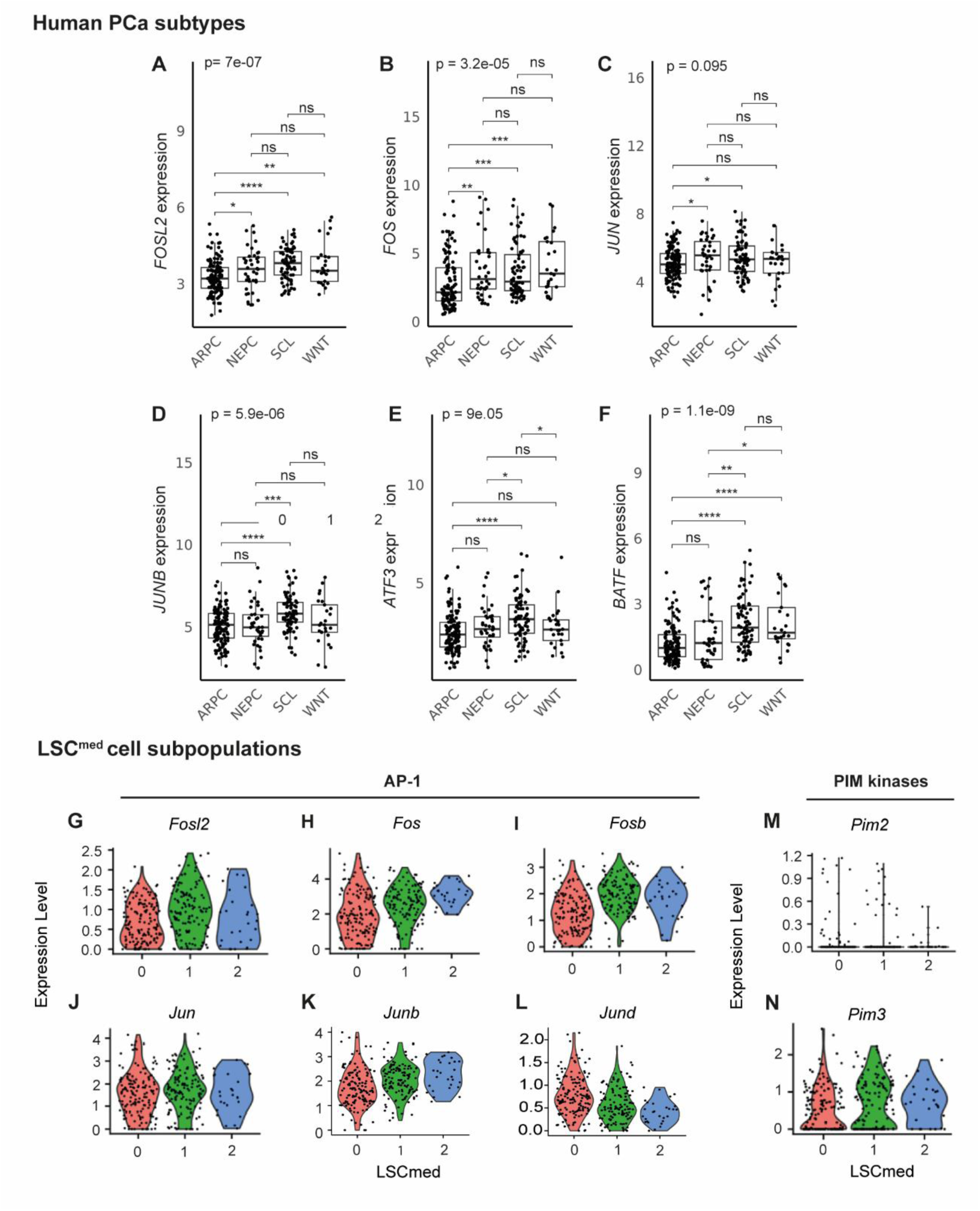
Expression of AP-1 family members and Pim kinases in human PCa subtypes and LSC^med^ cell subpopulations. **(A-F).** Expression of *FOSL2* (A), *FOS* (B), *JUN* (C), *JUNB* (D), *ATF3* (E) and *BATF* (F) in SU2C patient cohort (Data Ref: Abida *et al*., 2019a), according to the tumor molecular subtype (n=125 ARPC, n=40 NEPC, n=76 SCL, n=25 WNT). ns, not significant; *p<0.05; **p<0.01; ***p<0.001; ****p<0.0001 (Kruskal-Wallis with Holm-corrected Wilcoxon post-hoc tests). See Appendix Table S1 for exact p-values. (**G-N).** Expression of *Fosl2* (G), *Fos* (H), *Fosb* (I), *Jun* (J), *Junb* (K) and *Jund* (L), and of PIM kinases *Pim2* (**M**) and *Pim3* (**N**), in the three of LSC^med^ cell subpopulations.

**Fig EV5.**
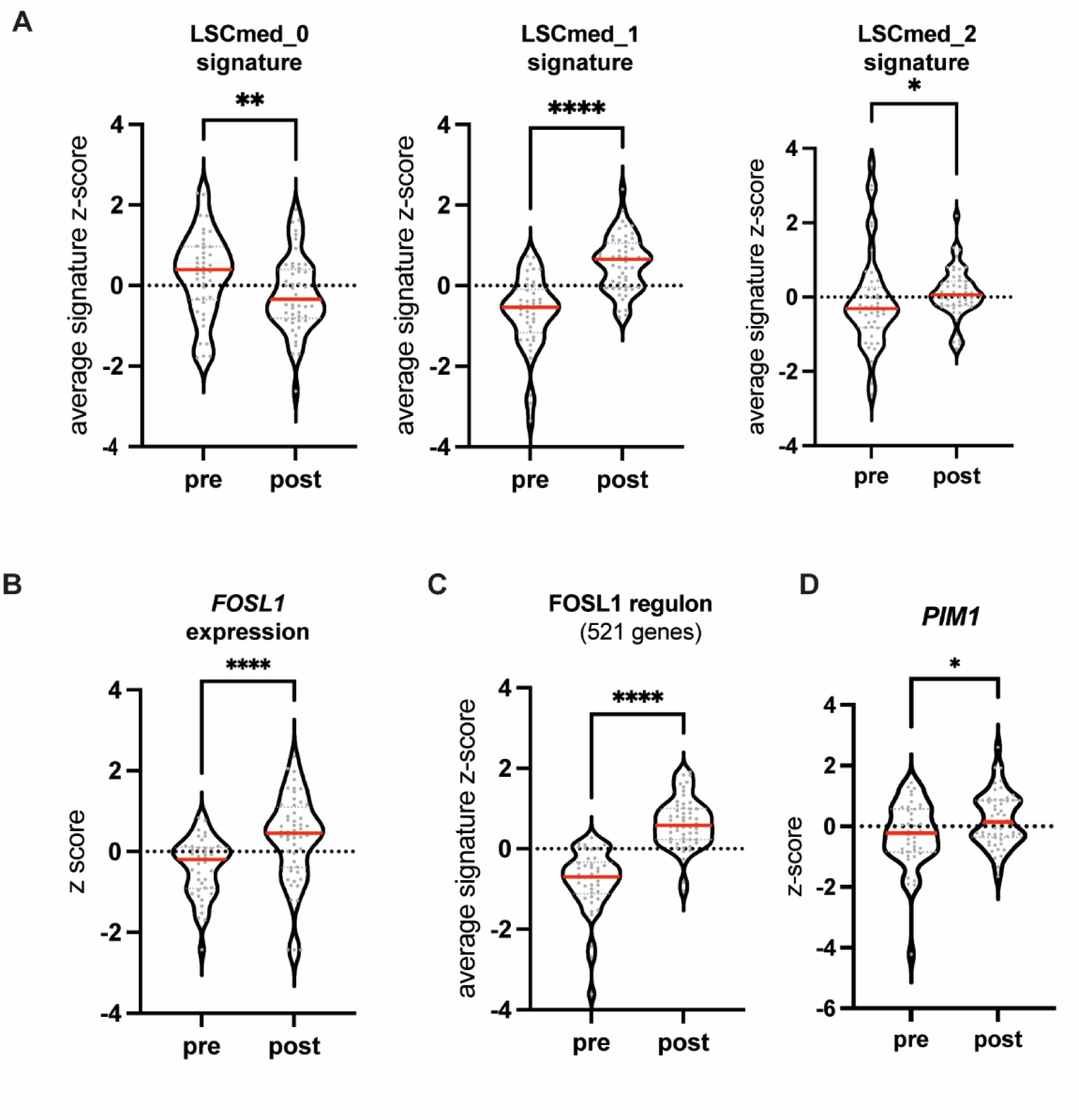
Induction of *FOSL1,* FOSL1 regulon, *PIM1* and LSC^med^ cluster 1 & 2 expression features upon enzalutamide treatment in human prostate cancer (DARANA study). RNA-sequencing data of naive (’pre’) and neoadjuvantly-treated (’post’) high-risk prostate cancer patients of the DARANA study (Data Ref: Linder *et al*., 2022b) was interrogated for LSC^med^ cluster signatures **(A)**, *FOSL1* (B), FOSL1 regulon **(C)** and *PIM1* (D) expression. Enzalutamide treatment induces *FOSL1,* FOSL1-regulon and LSC^med^-1 and LSC^med^ −2 gene expression and decreases LSC^med^-0 gene expression. *p<0.05; **p<0.01, ****p<0.0001 (Wilcoxon test). Exact p-values: p=0,0092 (LSC^med^-0), p=1,02e-40 (LSC^med^-1), p=0,0146 (LSC^med^-2), p=8,82e-18 (*FOSL1*), p=5,46e-58 (FOSL1 regulon), p=0,0378 (*PIM1*)

**Fig EV6.**
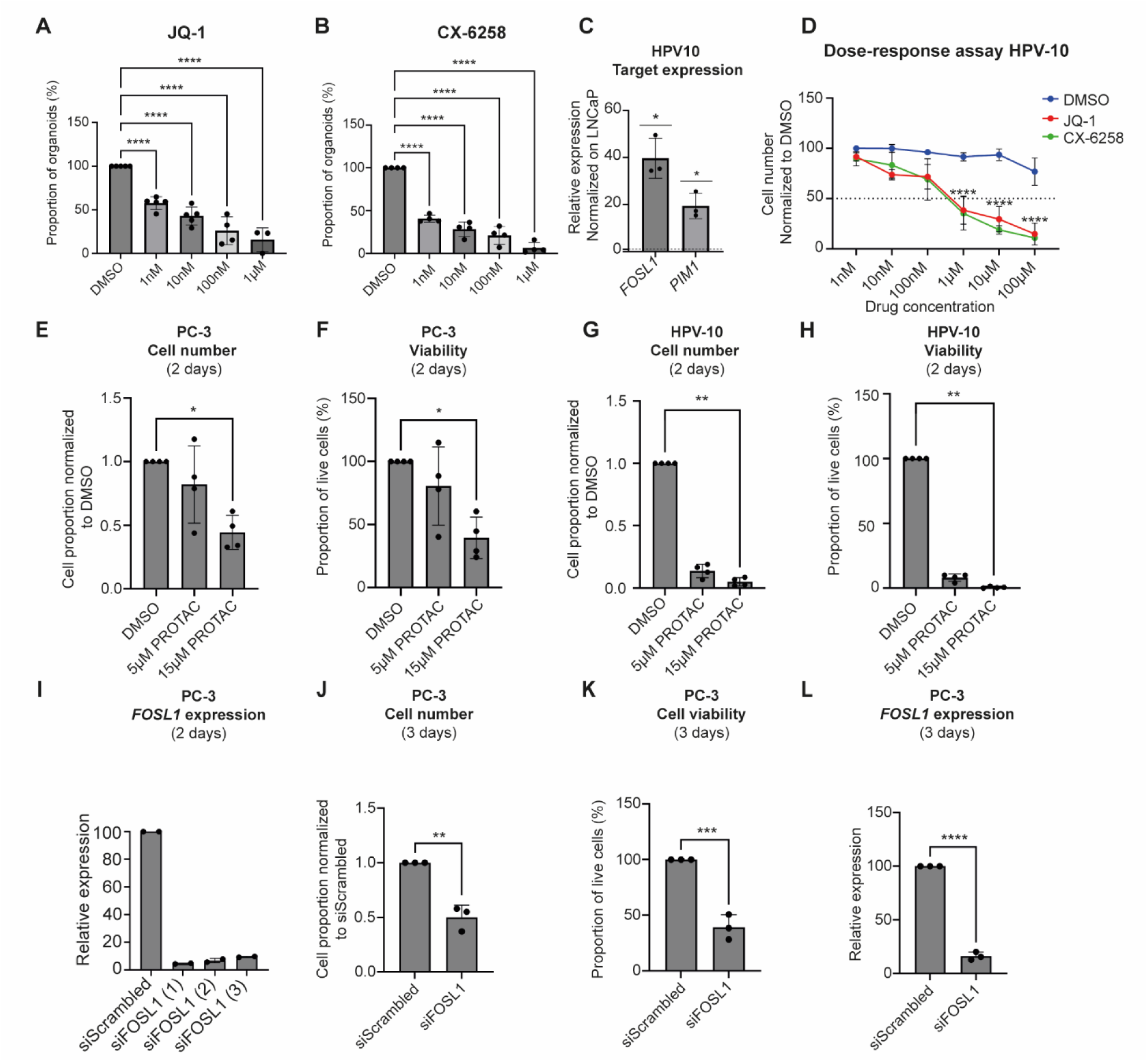
Targeting FOSL1 and PIM1 in Club-like cells *in vitro*. **(A,B)** Dose-response of JQ-1 **(A)** and CX-6258 **(B)** on the number of organoids formed by sorted *Pten^pc-/-^*LSC^med^ cells. Data are normalized to the DMSO condition (biological replicates, n=5 (A) and n=4 (B) independent experiments). ****p<0.0001 *versus* DMSO (ANOVA analysis and Dunnett’s *post hoc* test). Exact p-values are reported in Appendix Table S1. **(C)** Expression of *FOSL1* and *PIM1* in human HPV-10 cells determined by RT-qPCR. The results are normalized to the values obtained in LNCaP cells, represented by the horizontal dotted line (biological replicates, n = 3 independent experiments). *p<0.05 *versus* LnCAP cells (unpaired *t* test with Welch’s correction), p=0.0157 (*FOSL1*), p=0.0278 (*PIM1*). **(D)** Dose-response of JQ-1 and CX-6258 on the number of adherent HPV-10 cells. Data are normalized to the DMSO condition (each dot is the average of 3 biological replicates). ****p<0.0001 *versus* DMSO (ordinary two-way ANOVA with Šídák multiple comparisons test). Exact p-values are reported in Appendix Table S1. **(E-H)** Human PC-3 **(E,F)** and HPV-10 **(G,H)** cells were treated with T5224-PROTAC (’PROTAC’) at 0, 8, and 24h. The cells were collected at 48h. The cell number **(E,G)** and cell viability (adherent + in suspension) **(F,H)** were determined by trypan blue staining (biological replicates, n=4 independent experiments). The data are normalized to DMSO (control condition). *p<0.05, **p<0.01 (ANOVA analysis and Dunn’s post hoc test). **(I)** Human PC-3 cells were treated with siRNA (siScrambled or 3 different *FOSL1* siRNA, as indicated) for 6h and the cells were collected at 48h. The expression of *FOSL1* was measured by RT-qPCR (biological replicates, n=2 independent experiments). The data are normalized to siScrambled (control condition). The 3 siRNA *FOSL1* showed similar efficacy. ****p<0.0001 (ANOVA analysis and Dunnett’s post hoc test). Exact p-values are reported in Appendix Table S1. **(J-L)** Human PC-3 cells were treated with siScrambled or siFOSL1(1) for 6h and the cells were collected at 72h. Cell number **(J)** and cell viability (adherent + in suspension) **(K)** were determined by trypan blue staining. The expression of *FOSL1* **(L)** was measured by RT-qPCR. The data are normalized to siScrambled (biological replicates, n=3 independent experiments). **p<0.01, ***p<0.001, ****p<0.0001 (two-tailed t test). Exact p-values are reported in Appendix Table S1.

**Fig EV7.**
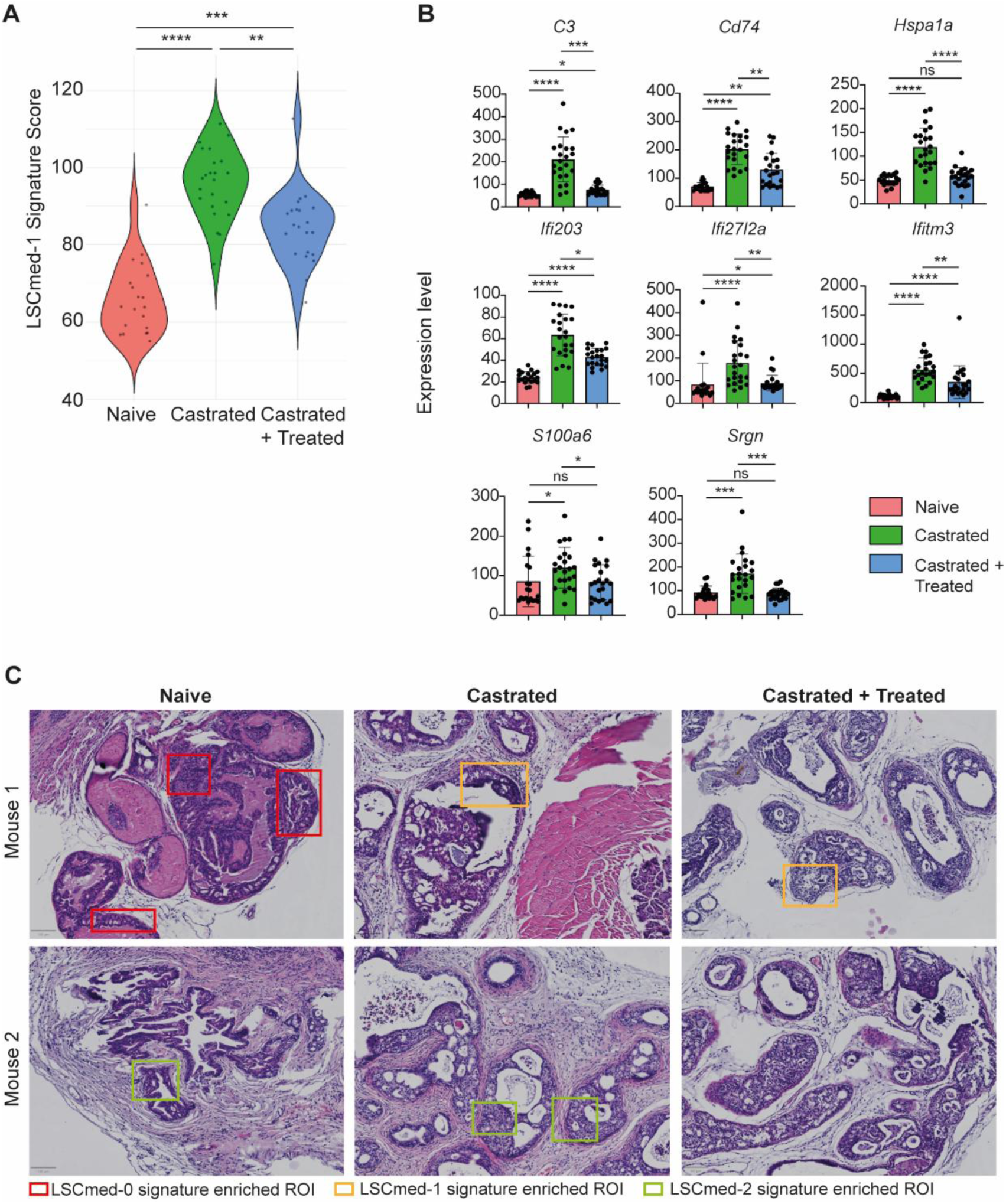
Digital spatial profiling of EPCAM+ cells in prostates from *Pten^pc^*^−/-^ mice non-castrated or castrated and treated with vehicle or JQ-1 and CX-6258. **(A)** Violin plot of the LSC^med^-1 signature score in selected ROIs of non-castrated (naïve), castrated vehicle-, or castrated JQ-1 and CX-6258-treated *Pten^pc-/-^* mice. n=10 – 13 ROI per mouse, 2 mice per condition. ns: not significant. *p< 0.05; ****p<0.0001 (non-parametric Kruskal-Wallis statistic tests). Exact p-values are reported in Appendix Table S1. **(B)** Transcript levels of the 8 most downregulated genes (p-value < 0.05 and a log2 fold change of <0.5) of the LSC^med^-1 and LSC^med^-2 cell signatures after the treatment in castrated mice. ns, not significant, *p<0.05, **p<0.001, ***p<0.001, **** p<0.0001 (Non-parametric Kruskal-Wallis and Dunnet post-hoc test). Exact p-values are reported in Appendix Table S1. **(C)** Representative H&E staining of prostates from naïve, castrated vehicle-, and castrated JQ-1 and CX-6258-treated *Pten^pc-/-^*mice. The colored squares represent LSC^med^-0, LSC^med^-1 and LSC^med^-2-enriched ROIs (color code indicated at the bottom of image). Scale bar: 100µm. n=2 mice per condition.

