## Supplemental data (Figures and Tables) for "Targeting Pre-Existing Club-Like Cells in Prostate Cancer Potentiates Androgen Deprivation Therapy"

*Baurès et al.*

Contents :

| | | |
|---|---|---|
| Appendix Figure S1 | FACS sorting of *Pten*$^{pc-/-}$ LSC$^{med}$ cells | Page 2 |
| Appendix Figure S2 | Organoid assay | Page 3 |
| Appendix Figure S3 | Time-course analysis of prostatic epithelial cell proliferation (Ki-67) and cell death (TUNEL) after castration. | Page 4 |
| Appendix Figure S4 | Molecular characterization of HPV-10 cells | Page 5 |
| Appendix Figure S5 | Assessment of the absence of drug toxicity in vivo. | Page 6 |
| Appendix Figure S6 | Combined JQ-1 and CX-6258 treatment reduces PC-3 cell growth in vivo | Page 7 |
| Appendix Figure S7 | Digital spatial profiling of EPCAM+ cells in prostates from *Pten*$^{pc-/-}$ mice non-castrated or castrated and treated with vehicle or JQ-1 and CX-6258 | Page 8 |
| Appendix Table S1 | Exact p-values | Page 9 |
| Appendix Table S2 | Histopathological analysis of kidney, lung and liver harvested from *Pten*$^{pc-/-}$ mice treated with vehicle *versus* JQ-1 + CX6258 | Page 12 |
| Appendix Table S3 | Primers used for RT-qPCR | Page 13 |

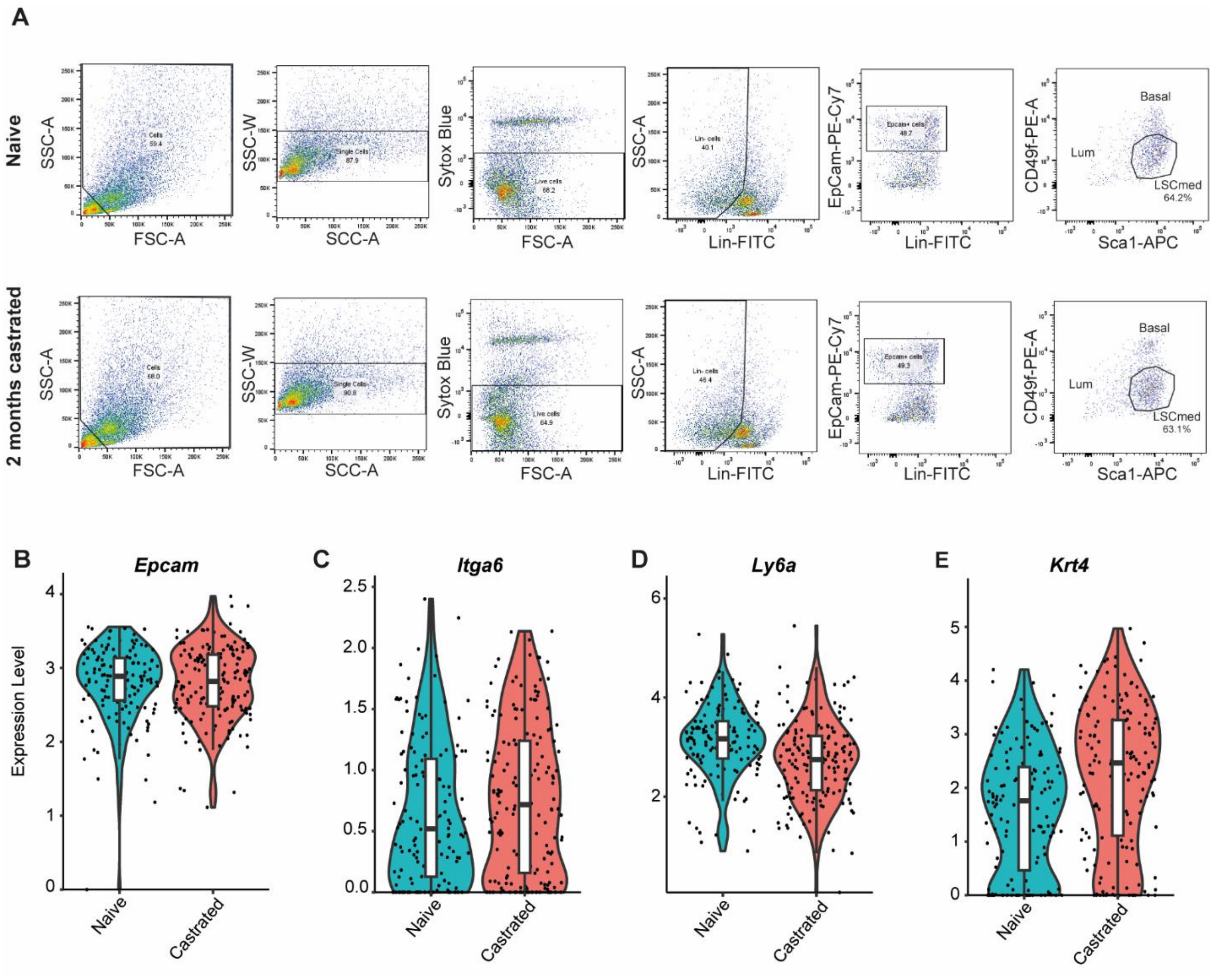

**Appendix Figure S1. FACS sorting of *Pten*$^{pc-/-}$ LSC$^{med}$ cells.**

**(A)** Representative experiment showing sequential gates used for the enrichment of LSC$^{med}$ cells from intact and castrated *Pten*$^{pc-/-}$ prostates. While LSC$^{med}$ cells account for ~80% of epithelial cells in *Pten*$^{pc-/-}$ mice irrespective of castration (Sackmann Sala et al, 2017), we used a more stringent gate for LSC$^{med}$ cell sorting to avoid any contamination with basal cells (hence <65% LSC$^{med}$, cells as indicated). See Sackmann Sala et al (2017) for additional information. **(B-E)** The identity of the cells analyzed by scRNAseq was assessed by the mRNA expression of the surface markers used in cell sorting: *Epcam* (B), *Sca-1/Ly6a* (C), *Cd49f/Itga6* (D) and the LSC$^{med}$ marker *Krt4* (E).

Figure complementary to Fig. 1, A to D.

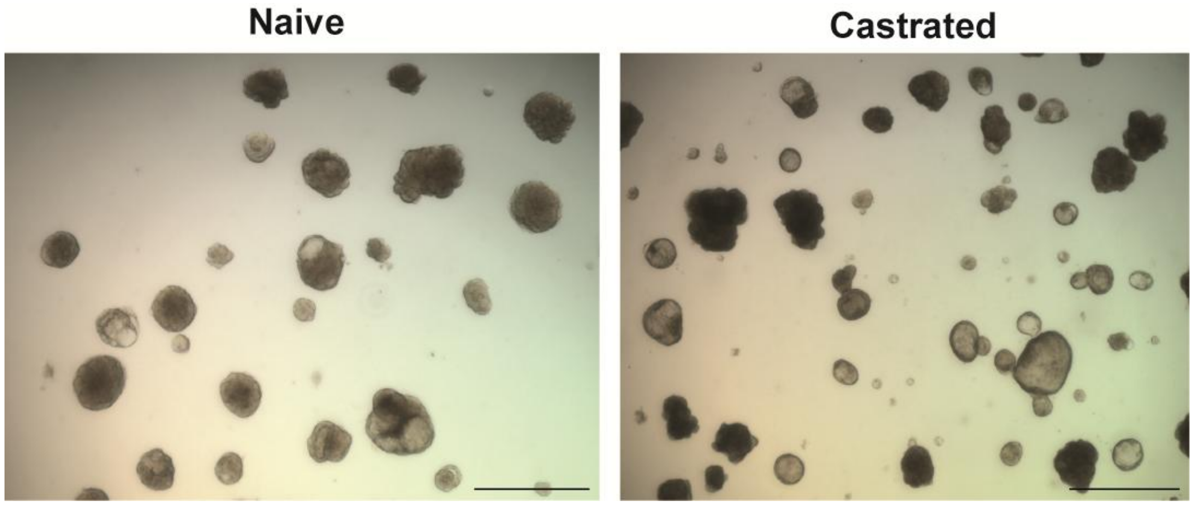

**Appendix Figure S2. Organoid assay.**

Representative images of organoids formed by $LSC^{med}$ cells sorted from intact *versus* 2-month-castrated $\textit{Pten}^{pc-/-}$ mice, as indicated. Scale bars: 100µm.

Figure complementary to Fig. 2K.

**A** Ki-67 cell counts

| | Naive | | Castrated 5 days | | Castrated 21 days | | Castrated 2 months | |
|---|---|---|---|---|---|---|---|---|
| | Positive | Total | Positive | Total | Positive | Total | Positive | Total |
| Mouse 1 | 2,296 | 15,602 | 1931 | 10,600 | 1244 | 10,838 | 954 | 11,711 |
| Mouse 2 | 1658 | 13,877 | 1190 | 9,156 | 1985 | 11,862 | 2765 | 9,533 |
| **Total** | **3954** | **29,479** | **3121** | **19,756** | **3229** | **22,700** | **3719** | **21,244** |

**B** TUNEL cell counts

| | Naive | | Castrated 5 days | | Castrated 21 days | | Castrated 2 months | |
|---|---|---|---|---|---|---|---|---|
| | Positive | Total | Positive | Total | Positive | Total | Positive | Total |
| Mouse 1 | 238 | 11,302 | 56 | 11,340 | 365 | 16,345 | 51 | 6,682 |
| Mouse 2 | 193 | 9,261 | 290 | 12,980 | 292 | 17,783 | 160 | 12,519 |
| Mouse 3 | 169 | 11,983 | / | / | 277 | 15,565 | 101 | 14,697 |
| Mouse 4 | 32 | 4,611 | / | / | / | / | 171 | 8,694 |
| **Total** | **632** | **37,157** | **346** | **24,320** | **934** | **49,693** | **483** | **42,592** |

**C** TUNEL assay

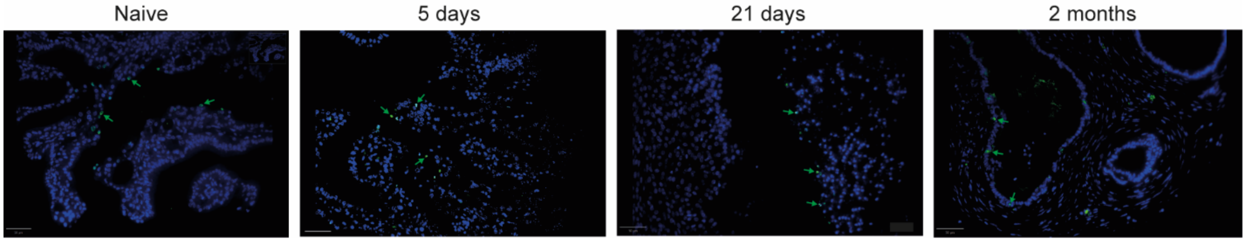

**Appendix Figure S3. Time-course analysis of prostatic epithelial cell proliferation (Ki-67) and cell death (TUNEL) after castration.**

The effect of castration on prostatic epithelial cell proliferation and cell death was determined by Ki-67 immunohistochemistry and TUNEL assay, respectively, in prostates from 2 to 4 intact *Pten*$^{pc-/-}$ mice *versus* 5 days, 21 days and 2 months after castration. The number of positive and total cell counted per condition is reported in **A** (Ki-67) and **B** (TUNEL). Representative images of prostate sections after the TUNEL assay for each time point are shown in **C** (scale bar: 250 µm).

Figure complementary to Fig. 3, C and D

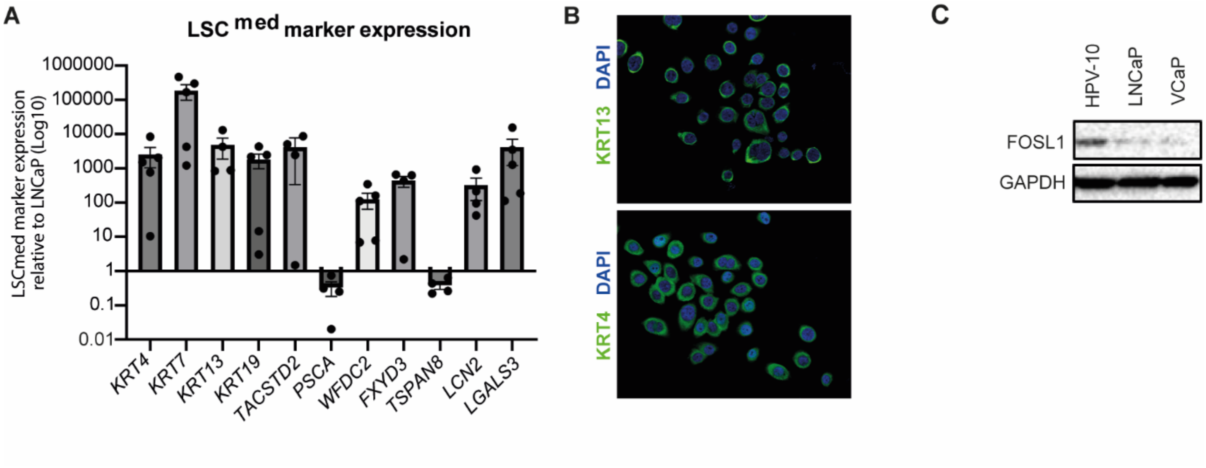

**Appendix Figure S4. Molecular characterization of HPV-10 cells.**

**(A)** Expression of LSC$^{med}$ markers in human HPV-10 cells as determined by RT-qPCR. The results are normalized to the values obtained in LNCaP cells, represented by the horizontal dotted line. n=4 to 5 independent experiments performed in duplicate. **(B)** Immunofluorescence analysis of KRT4 and KRT13 in HPV-10 cells. **(C)** Immunoblot showing the level of FOSL1 expression in HPV-10 compared to two luminal cell lines (LNCaP, VCaP).

Figure complementary to Fig. 5H-J.

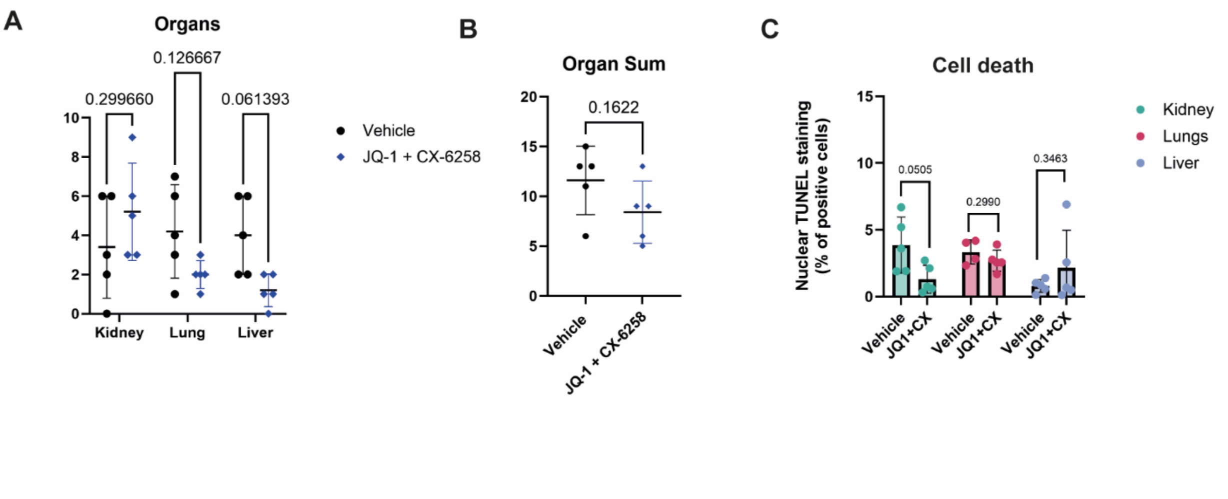

**Appendix Figure S5. Assessment of the absence of drug toxicity *in vivo*.**

**(A,B)** Histological analysis of kidneys, lungs, and livers harvested from five *Pten*$^{pc-/-}$ mice in both the vehicle- and JQ-1/CX-6258-treated groups was performed, and degenerative and inflammatory lesions were rated on a 5-point scale (0 = no lesion, 1 = scattered, 2 = mild, 3 = moderate, 4 = marked), as reported in Appendix Table S2. Scores for each animal per organ (A) and all organs combined (B) are shown. Data were compared by Unpaired *t*-test (organ sum) and multiple unpaired *t*-tests (individual organs). **(C)** The 3 tissues from the same animals were analyzed by the TUNEL assay to quantify cell death. Unpaired *t*-test with Welch's correction.

Figure complementary to Fig. 6, A-F.

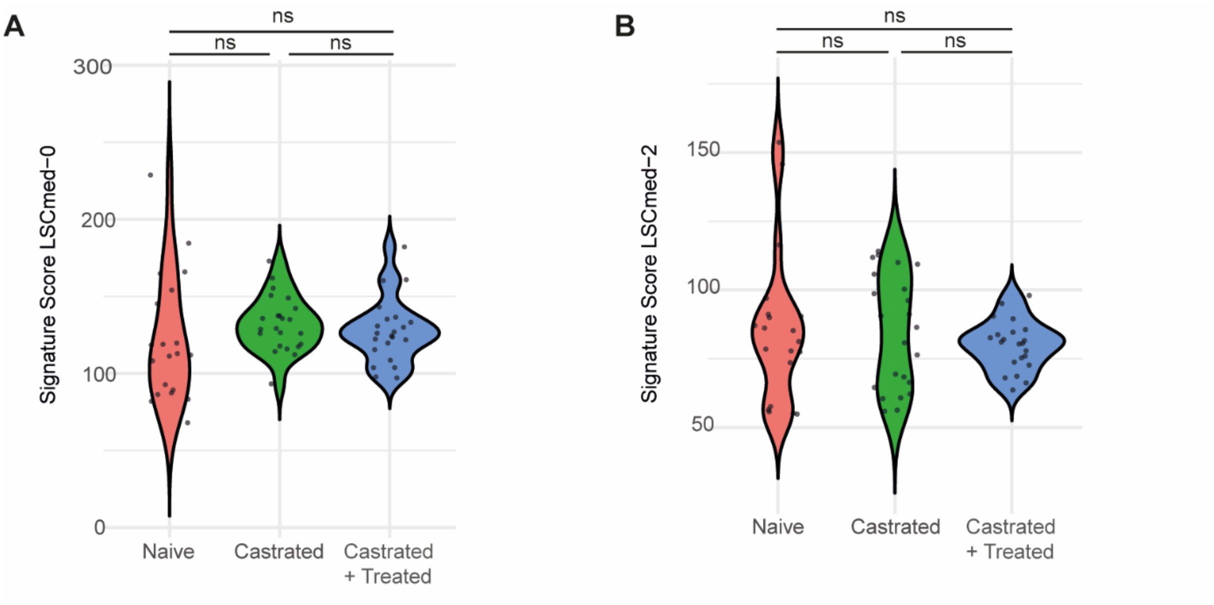

**Appendix Figure S6. Digital spatial profiling of EPCAM+ cells in prostates from *Pten*$^{pc-/-}$ mice non-castrated or castrated and treated with vehicle or JQ-1 and CX-6258.**

Violin plot of the LSC$^{med}$-0 **(A)** and LSC$^{med}$-2 **(B)** signature score in selected ROIs of non-castrated (naive), castrated vehicle-, or castrated + JQ-1/CX-6258-treated *Pten*$^{pc-/-}$ mice. n=10 to 13 ROI per mouse, 2 mice per condition. ns: not significant by non-parametric Kruskal-Wallis statistical tests. Exact p values are reported in Appendix Table S1.

Figure complementary to Figure EV7.

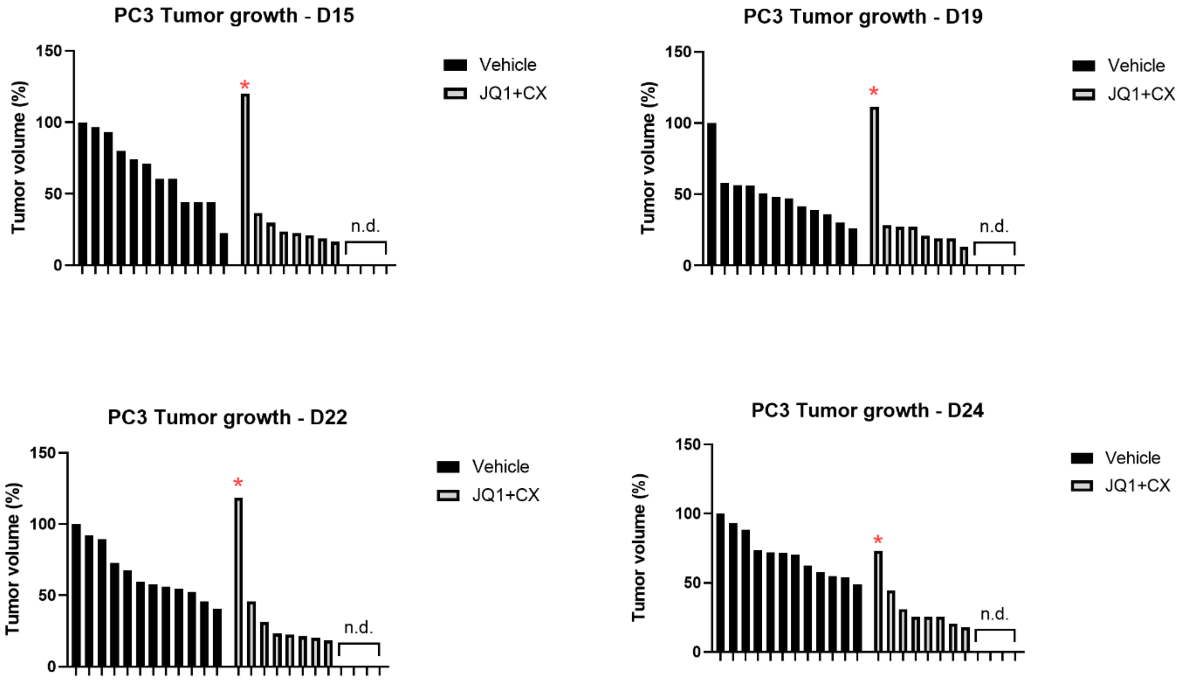

**Appendix Figure S7. Combined JQ-1 and CX-6258 treatment reduces PC-3 cell growth *in vivo***

The size of individual tumors grown from PC-3 cells xenografted into NSG mice is shown at days 15, 19, 22 and 24 after treatment. Before tumors were measurable, two mice (i.e. 4 xenografts) in the treated group experienced issues with the gavage procedure that prevented continuation of oral treatment for ethical reasons (n.d.). One tumor in the treated group initially grew abnormally fast (*) and regressed at the last timepoint, therefore it was not included in the analysis. The tumor on the other flank of the same mouse behaved normally.

Figure complementary to Fig. 6, G-J.

### Appendix Table S1. Exact p-values

#### Fig 3, E to G

| Fig 3E | | | Fig 3F | | | Fig 3G | | |
|---|---|---|---|---|---|---|---|---|
| *Gene* | Naive vs 5 days | Naive vs 21 days | *Gene* | Naive vs 5 days | Naive vs 21 days | *Gene* | Naive vs 5 days | Naive vs 21 days |
| *Oit1* | 0,6272 | 0,0003 | *Ly6d* | 0,2125 | 0,2362 | *Muc4* | 0,1531 | 0,0382 |
| *Ly6c1* | 0,3756 | 0,0625 | *Cd44* | 0,932 | 0,0816 | *Lgals4* | 0,7301 | 0,8479 |
| *Hoxb13* | 0,1461 | 0,0003 | *Bcar1* | 0,3855 | 0,1178 | *Hnf4a* | 0,3464 | 0,7893 |
| *Spink5* | 0,0029 | 0,0004 | *Fosl1* | 0,252 | 0,1102 | *Onecut2* | 0,318 | 0,599 |
| | | | *Pim1* | 0,2778 | 0,2619 | *Ugt2b34* | 0,392 | 0,3887 |
| | | | *F3* | 0,1865 | 0,1682 | *Reg1* | 0,3433 | 0,1483 |
| | | | *Tm4sf1* | 0,3356 | 0,1929 | | | |
| | | | *Ifi202b* | 0,3792 | 0,1783 | | | |

#### Fig 4, I and J

| PCa subtypes | Fig 4I | Fig 4J |
|---|---|---|
| | *FOSL1* | FOSL1 regulon |
| ARPC vs NEPC | 0.27 | 0.0013 |
| ARPC vs SCL | 1.7e-14 | 2.22e-16 |
| ARPC vs WNT | 0.00021 | 2.2e-05 |
| NEPC vs SCL | 2.7e-06 | 2e-05 |
| NEPC vs WNT | 0.014 | 0.056 |
| SCL vs WNT | 0.24 | 0.045 |

#### Figure 5

| | Fig 5C | Fig 5D | Fig 5H | Fig 5J | Fig 5K | Fig 5L | Fig 5M |
|---|---|---|---|---|---|---|---|
| JQ-1 | 0,0007 | 0,0037 | 0,0015 | 0,3275 | 0,0099 | 0,8159 | 0,0005 |
| CX-6258 | 0,0007 | 0,0053 | 0,0012 | 0,1203 | 0,0059 | 0,4861 | 0,0008 |
| JQ-1 + CX-6258 | <0,0001 | 0,0003 | 0,0002 | 0,0098 | 0,0003 | 0,012 | 0,0002 |

### Fig EV1

| Fig EV1A | | Fig EV1B | | Fig EV1C | |
|---|---|---|---|---|---|
| *Gene* | LSCmed-0 markers | *Gene* | LSCmed-1 markers | *Gene* | LSCmed-2 markers |
| *Oit1* | 0.0754 | *Bcar1* | 0.0247 | *Ugt2b34* | 0.1511 |
| *Hoxb13* | 0.2050 | *Ly6d* | 0.1207 | *Hnf4a* | 0.4917 |
| *Ly6c1* | 0.0038 | *Cd44* | 0.1953 | *Muc4* | 0.2525 |
| *Spink5* | 0.0001 | *Fosl1* | 0.0383 | *Onecut2* | 0.4501 |
| | | *Pim1* | 0.0476 | *Reg1* | 0.1327 |
| | | *F3* | 0.0129 | *Ugt2b34* | 0.1511 |
| | | *Tm4sf1* | 0.0650 | | |
| | | *Ifi202b* | 0.3704 | | |

### Fig EV4, A to F

| | Fig EV4A | Fig EV4B | Fig EV4C | Fig EV4D | Fig EV4E | Fig EV4F |
|---|---|---|---|---|---|---|
| | *FOSL2* | *FOS* | *JUN* | *JUNB* | *ATF3* | *BATF* |
| ARPC vs NEPC | 0.031 | 0.003 | 0.039 | 0.9 | 0.086 | 0.36 |
| ARPC vs SCL | 5.7e-08 | 0.00011 | 0.046 | 2.8e-07 | 6.1e-06 | 3.4e-10 |
| ARPC vs WNT | 0.0036 | 0.00099 | 0.66 | 0.31 | 0.23 | 5.7e-05 |
| NEPC vs SCL | 0.096 | 0.93 | 0.75 | 0.0007 | 0.049 | 0.0033 |
| NEPC vs WNT | 0.49 | 0.37 | 0.29 | 0.45 | 0.86 | 0.016 |
| SCL vs WNT | 0.58 | 0.36 | 0.42 | 0.069 | 0.047 | 0.88 |

### Figure EV6

| vs DMSO | Fig EV6 A (JQ-1) | Fig EV6 B (CX6258) | Fig EV6 D (JQ-1) | Fig EV6D (CX-6258) |
|---|---|---|---|---|
| 1 nM | <0.0001 | <0.0001 | 0,9990 | 0,9885 |
| 10 nM | <0.0001 | <0.0001 | 0,0539 | 0,6048 |
| 100 nM | <0.0001 | <0.0001 | 0,0979 | 0,0424 |
| 1 µM | <0.0001 | <0.0001 | <0,0001 | <0,0001 |
| 10µM | / | / | <0,0001 | <0,0001 |
| 100µM | / | / | <0,0001 | <0,0001 |

| vs DMSO | Fig EV6 E (PC-3 / PROTAC) | Fig EV6 F (PC-3 / PROTAC) | Fig EV6 G (HPV-10 / PROTAC) | Fig EV6 H (HPV-10 / PROTAC) |
|---|---|---|---|---|
| 5 µM | 0.9546 | 0.6364 | 0.2171 | 0.3307 |
| 15 µM | 0.0284 | 0.0189 | 0.0082 | 0.0042 |

| vs siScrambled | Fig EV6 I (siFosl1) | Fig EV6 J (siFosl1) | Fig EV6 K (siFosl1) | Fig EV6 L (siFosl1) |
|---|---|---|---|---|
| Fosl1(1) | <0.0001 | 0.0016 | 0.0007 | <0.0001 |
| Fosl1(2) | <0.0001 | NA | NA | NA |
| Fosl1(3) | <0.0001 | NA | NA | NA |

*NA=Not applicable*

### Fig EV7A

| Score | Naive vs castrated | Naive vs castrated + treated | Castrated vs castrated + treated |
|---|---|---|---|
| LSCmed-1 | 4.60e-10 | 0.00033 | 0.0062 |

### Fig EV7B

| *Gene* | Naive vs castrated | Naive vs castrated + treated | Castrated vs castrated + treated |
|---|---|---|---|
| C3 | 8.02255e-10 | 0.01407 | 0.00012 |
| Cd74 | 6.39255e-10 | 0.00149 | 0.00192 |
| Hspa1a | 2.88358e-08 | 0.262841 | 3.49953e-06 |
| Ifi203 | 6.87031e-11 | 3.837772e-05 | 0.01263 |
| Ifi27l2a | 3.20403e-07 | 0.01797 | 0.00408 |
| Ifitm3 | 3.50697e-11 | 5.97560e-05 | 0.00695 |
| S100a6 | 0.04041 | 0.77878 | 0.04041 |
| Srgn | 0.00053 | 0.72156 | 0.00016 |

### Appendix Fig S7

| Score | Naive vs castrated | Naive vs castrated + treated | Castrated vs castrated + treated |
|---|---|---|---|
| LSCmed-0 | 0.1065 | 0.2959 | 0.4123 |
| LSCmed-2 | 0.6978 | 0.6978 | 0.6978 |

**Appendix Table S2. Histopathological analysis of kidney, lung and liver harvested from *Pten*$^{pc-/-}$ mice treated with vehicle *versus* JQ-1 + CX-6258**

Complementary to Appendix Figure S5

Occasional = 1
Low grade = 2
Medium grade = 3
High grade = 4

| | | KIDNEY | | | | | | | LUNG | | | | | | | | LIVER | | | | | | |
|---|---|---|---|---|---|---|---|---|---|---|---|---|---|---|---|---|---|---|---|---|---|---|---|
| Mouse ID | Treatment | High grade cytoplasmic vacuolation | Inflammation | Tubular regeneration | Protein tubular casts | Calcification | Sclerosed glomerulus | Sum | Emphysema | Edema | Inflammation | Erythrocytes | Macrophages | Pleuritis | BALT hyperplasia | Sum | Microgranulomas | Lipidosis | Kupffer proliferation | Inflammation | Cell edema | Sum | Organ Sum |
| 347 | Vehicle | 0 | 0 | 0 | 0 | 0 | 0 | 0 | 2 | 2 | 0 | 0 | 0 | 0 | 0 | 4 | 2 | 0 | 0 | 0 | 0 | 2 | 6 |
| 329 | Vehicle | 0 | 1 | 1 | 0 | 0 | 0 | 2 | 2 | 0 | 2 | 2 | 1 | 0 | 0 | 7 | 1 | 0 | 0 | 0 | 1 | 2 | 11 |
| 328 | Vehicle | 1 | 2 | 0 | 0 | 0 | 0 | 3 | 0 | 0 | 3 | 0 | 2 | 1 | 0 | 6 | 2 | 1 | 1 | 0 | 0 | 4 | 13 |
| 278 | Vehicle | 1 | 2 | 1 | 2 | 0 | 0 | 6 | 0 | 2 | 0 | 0 | 0 | 1 | 0 | 3 | 2 | 2 | 0 | 2 | 0 | 6 | 15 |
| 277 | Vehicle | 1 | 0 | 1 | 3 | 1 | 0 | 6 | 0 | 0 | 0 | 0 | 1 | 0 | 0 | 1 | 2 | 2 | 0 | 2 | 0 | 6 | 13 |
| 1634 | JQ-1 / CX6258 | 0 | 0 | 1 | 2 | 0 | 0 | 3 | 0 | 0 | 0 | 0 | 2 | 0 | 1 | 3 | 0 | 0 | 0 | 0 | 0 | 0 | 6 |
| 1632 | JQ-1 / CX6259 | 0 | 0 | 1 | 2 | 0 | 0 | 3 | 0 | 1 | 0 | 0 | 0 | 0 | 0 | 1 | 0 | 0 | 0 | 0 | 1 | 1 | 5 |
| 364 | JQ-1 / CX6260 | 0 | 0 | 2 | 3 | 0 | 0 | 5 | 0 | 1 | 1 | 0 | 0 | 0 | 0 | 2 | 0 | 2 | 0 | 0 | 0 | 2 | 9 |
| 316 | JQ-1 / CX6261 | 0 | 0 | 2 | 2 | 1 | 1 | 6 | 0 | 2 | 0 | 0 | 0 | 0 | 0 | 2 | 0 | 1 | 0 | 0 | 0 | 1 | 9 |
| 313 | JQ-1 / CX6262 | 0 | 3 | 1 | 3 | 1 | 1 | 9 | 0 | 2 | 0 | 0 | 0 | 0 | 0 | 2 | 1 | 1 | 0 | 0 | 0 | 2 | 13 |

### Appendix Table S3. Primers used for RT-qPCR

| Target gene | Species | Forward / Reverse | Sequences |
|---|---|---|---|
| Cyclophilin-A | Mouse | F | CAG-GTC-CTG-GCA-TCT-TGT-CC |
| | | R | TTG-CTG-GTC-TTG-CCA-TTC-CT |
| Oit1 | Mouse | F | GAT-GAA-AAA-GGA-CAG-CTT-TG |
| | | R | TTT-GTC-GTT-CAT-TTT-GGT-CC |
| Hoxb13 | Mouse | F | ATG-CAG-CCA-ACA-AGT-TTA-TC |
| | | R | CAA-GAA-CCT-TCT-TCT-CCT-TG |
| Ly6c1 | Mouse | F | GAT-TGA-GGA-CTC-TCA-AAG-AAG |
| | | R | ATG-TTA-GGA-TCC-CTG-ATT-GG |
| Spink5 | Mouse | F | GTG-CTG-AGA-TAT-TCA-TGA-GG |
| | | R | CTT-TCT-TCA-GCA-TCT-TTC-TGG |
| Bcar1 | Mouse | F | GTA-TGG-CCA-GGA-GGT-ATA-TG |
| | | R | GTC-ATA-CAC-ATC-CAA-CAA-TGG |
| F3 | Mouse | F | CCC-CAA-AGT-TTT-TAC-CTT-ACC |
| | | R | TAT-ATA-ACC-CAA-GTC-CTT-GCC |
| Fosl1 | Mouse | F | AGC-AGA-AGT-TCC-ACC-TTG |
| | | R | CTA-GGG-CTC-GTA-TGA-CTC |
| Ly6D | Mouse | F | CAT-ATA-TAG-CAC-AGT-GCC-TC |
| | | R | CAA-GTA-CAA-AAT-CAG-AGG-GG |
| Tm4sf1 | Mouse | F | CTC-GCC-AAC-AGC-AAT-ATA-AC |
| | | R | GCA-CAT-TTC-CTT-ACA-AAA-GC |
| Pim1 | Mouse | F | CAA-ACT-GTC-TCT-TCA-GAG-TG |
| | | R | GTT-CCG-GAT-TTC-TTC-AAA-GG |
| Ifi202b | Mouse | F | TTT-CAT-CAA-GGG-AGA-AAA-GC |
| | | R | GTT-TTG-TTG-CTT-TCA-ATG-CC |
| CD44 | Mouse | F | GAA-TTA-GCT-GGA-CAC-TCA-AG |
| | | R | CAC-CTT-CTC-CTA-CTA-TTG-ACC |
| HNF4a | Mouse | F | TCC-TAG-GCA-ATG-ACT-ACA-TC |
| | | R | CTG-GAT-CAA-AGA-AGA-TGA-TGG |
| Reg1 | Mouse | F | CAT-GAT-CCC-AAA-AGG-AAT-CG |
| | | R | GTT-TGA-AGT-CAG-AGA-TAC-ACA-G |
| Ugt2b34 | Mouse | F | GGG-TAG-CAT-AAC-AGA-AGA-AAG |
| | | R | GGT-TTT-GGA-ATG-ACC-TAG-AAG |
| Onecut2 | Mouse | F | CTG-GAG-TAA-ACT-CAA-ATC-TGG |
| | | R | TTC-CTG-TCT-TTG-TTT-GGT-TC |
| Lgals4 | Mouse | F | TTC-TTT-GAT-CTG-TCA-ATC-CG |
| | | R | GTG-ATG-TCA-CCA-TTG-ATC-TC |

| | | | |
|---|---|---|---|
| Muc4 | Mouse | F | GTC-AAT-GTT-CCT-GCC-TAT-AC |
| | | R | CAA-AAT-ACC-CAT-CTC-CAC-TG |
| Actin B | Human | F | AAG-ACC-TGT-ACG-CCA-ACA-CA |
| | | R | TGA-TCT-CCT-TCT-GCA-TCC-TG |
| Fosl1 | Human | F | CTT-GTG-AAC-AGA-TCA-GCC |
| | | R | CCA-GAT-TTC-TCA-TCT-TCC-AG |
| Pim1 | Human | F | CTC-TTC-AGA-ATG-TCA-GCA-TC |
| | | R | GGA-TGG-TTC-TGG-ATT-TCT-TC |
| Krt4 | Human | F | GTA-CAG-AAT-GTC-TGG-AGA-ATG |
| | | R | TGG-TAG-AGA-TGA-TCT-TGC-TG |
| Krt7 | Human | F | CTC-TGT-GAT-GAA-TTC-CAC-TG |
| | | R | ATG-GAA-TAA-GCC-TTC-AGG-AG |
| Krt13 | Human | F | ATA-CGC-TTT-GGT-TTC-TCA-AC |
| | | R | TTA-TGT-TTG-CAG-AAA-GGC-AG |
| Krt19 | Human | F | AAC-CAT-GAG-GAG-GAA-ATC-AG |
| | | R | CAT-GAC-CTC-ATA-TTG-GCT-TC |
| Tacstd2 | Human | F | GCA-GAA-CAC-GTC-TCA-GAA |
| | | R | TTG-ATG-TCC-CTC-TCG-AAG-TA |
| PSCA | Human | F | AAG-TGG-ACT-GAG-TAG-AAC-TG |
| | | R | GTG-TTT-ATT-AAG-GGC-CTA-CG |
| Wfdc2 | Human | F | AAT-GAT-AAG-GAG-GGT-TCC-TG |
| | | R | GAA-ACT-TTC-TCT-CCT-CAC-TG |
| Fxyd3 | Human | F | CTT-CTG-CTG-ATC-CTG-AAA-TTG |
| | | R | CCT-CAC-TTC-TTT-TCC-TTA-GAT-G |
| Tspan8 | Human | F | CTA-AGT-CTG-ATC-GCA-TTG-TG |
| | | R | AAC-ACA-ATT-ATG-GCT-TCC-TG |
| Lcn2 | Human | F | GGA-AAA-AGA-AGT-GTG-ACT-ACT-G |
| | | R | GTA-ACT-CTT-AAT-GTT-GCC-CAG |
| Lgals3 | Human | F | AGA-TTT-CCA-AAG-AGG-GAA-TG |
| | | R | AAG-TGC-AAA-CAA-TGA-CTC-TC |
